# Adrenomedullin 2/intermedin is a slow off-rate, long-acting endogenous agonist of the adrenomedullin_2_ G protein-coupled receptor

**DOI:** 10.1101/2023.01.13.523955

**Authors:** Katie M. Babin, Jordan A. Karim, Peyton H. Gordon, James Lennon, Alex Dickson, Augen A. Pioszak

## Abstract

The signaling peptides adrenomedullin 2/intermedin (AM2/IMD), adrenomedullin (AM), and CGRP have overlapping and distinct functions in the cardiovascular, lymphatic, and nervous systems by activating three shared receptors comprised of the class B GPCR CLR in complex with a RAMP1, −2, or −3 modulatory subunit. Here, we report that AM2/IMD, which is thought to be a non-selective agonist, is kinetically selective for CLR-RAMP3, known as the AM_2_R. AM2/IMD-AM_2_R elicited substantially longer duration cAMP signaling than the eight other peptide-receptor combinations due to AM2/IMD slow off-rate binding kinetics. The regions responsible for the slow off-rate were mapped to the AM2/IMD mid-region and the RAMP3 extracellular domain. MD simulations revealed how these bestow enhanced stability to the complex. Our results uncover AM2/IMD-AM_2_R as a cognate pair with unique temporal features, define the mechanism of kinetic selectivity, and explain how AM2/IMD and RAMP3 collaborate to shape the signaling output of a clinically important GPCR.

## Introduction

Adrenomedullin 2/intermedin (AM2/IMD) was discovered by two groups in 2004 based on its similarity to the calcitonin gene-related peptide (CGRP) family of peptides, which also includes adrenomedullin (AM) and amylin^1,2^. AM2/IMD has overlapping and distinct functions, but it is the least understood of these peptides. Like CGRP and AM, when administered peripherally AM2/IMD induces vasodilation, and it has protective effects in the cardiovascular, pulmonary, and renal systems^3^. AM2/IMD stabilizes the endothelial barrier^4^, has anti-inflammatory actions^5^, and protects against sepsis in mice^6^. AM2/IMD is angiogenic^7^ and its knock-out in mice revealed roles in enlarging the vascular lumen and promoting vessel fusion^8,9^. Beyond the circulatory system, AM2/IMD protected against obesity and insulin resistance in mice by promoting beige cell biogenesis and reducing adipose inflammation^10–12^, and its central administration activated the hypothalamic-pituitary-adrenal axis and increased sympathetic activity^13,14^. There is promise in targeting AM2/IMD signaling for therapeutics for cardiovascular, metabolic and other disorders^15^, but progress has been hindered by our limited understanding of the mechanisms that distinguish AM2/IMD signaling from that of AM and CGRP.

AM2/IMD, AM, and CGRP share three heterodimeric receptors that are comprised of a common class B G protein-coupled receptor (GPCR) subunit, the calcitonin receptor-like receptor (CLR), and a variable receptor activity modifying protein (RAMP1-3) subunit that alters CLR ligand selectivity^16^. These couple most efficiently to the Gs protein to activate cAMP signaling, but they also couple to Gq and Gi proteins^17,18^. Much of our understanding of the pharmacology of the three peptides and receptors comes from studies of cAMP signaling^19^. CGRP is most potent at the CLR-RAMP1 complex, which is designated the CGRP receptor. This mediates CGRP actions in the trigeminovascular system and is the target of several recently approved inhibitor drugs for migraine headache^20^. AM is most active and equally potent at the CLR-RAMP2 and CLR-RAMP3 complexes, which are termed the AM_1_ and AM_2_ receptors, respectively. The AM_1_ receptor mediates the essential developmental actions of AM in the cardiovascular and lymphatic systems^21–26^. Why cells need a second AM receptor is unclear as AM_2_R functions remain poorly defined.

AM2/IMD is thought to be relatively non-selective for the three CLR-RAMP complexes^1^. It exhibits a slight preference for the AM_2_R, but the AM_2_ nomenclature is not meant to imply that it is the AM2/IMD receptor^19,27^. In cAMP assays, AM2/IMD is slightly less potent than CGRP and AM at the CGRP and AM_1_ receptors, respectively, and is equipotent to AM at the AM_2_R. Different effects of AM2/IMD have been ascribed to signaling through either the CGRP or the AM receptors, but in many cases, the receptor(s) mediating a given AM2/IMD action is unclear. There are examples of apparent mismatch between AM2/IMD pharmacology at the cloned receptors and *in vivo*, but the bases for these discrepancies are unknown^27^. Recently, the idea that AM2/IMD, AM, and CGRP promote distinct receptor-transducer coupling profiles (agonist bias) has been explored. There are some data supporting this^18,28^, but other studies found less evidence for bias^17,29^, so the extent to which signaling bias distinguishes the peptides remains unclear.

Structural studies advanced our understanding of how the peptides bind the receptors and how the RAMPs modulate binding. In crystal structures of CLR-RAMP1/2-peptide ECD complexes, the C-terminal fragments of CGRP, AM, and AM2/IMD occupied a shared binding site on the CLR ECD, but with distinct, mostly unstructured conformations^30,31^. RAMP1/2 augmented the ECD binding site with unique contacts to the peptides that contributed to ligand selectivity^30,32,33^. Cryo-EM structures of several peptide-bound CLR-RAMP complexes with Gs showed that the N-terminal half of each peptide adopted a disulfide loop and α-helix structure that similarly occupied the CLR TMD^34,35^. 3D variability analyses of the cryo-EM data were consistent with differential RAMP1-3 modulation of CLR ECD-TMD inter-domain dynamics playing a role in receptor phenotype^34^. Despite this progress, several issues remain unresolved. How the two AM peptides bind the CLR-RAMP3 ECD complex is unclear because their C-terminal fragments in the AM_2_R cryo-EM structures were modeled with different conformations than in the crystal structures. The relative contributions of RAMP augmentation of the CLR ECD peptide binding site and modulation of CLR dynamics to receptor phenotype is unclear. In addition, it is unknown if the peptide agonists differentially modulate CLR dynamics and/or work in concert with the RAMPs to shape the signaling outcomes.

The temporal features of AM2/IMD, AM, and CGRP signaling have received little attention. Here, we test the hypothesis that differences in their receptor binding and signaling kinetics distinguish their signaling capabilities. Strikingly, we find that AM2/IMD exhibits significantly longer duration cAMP signaling at the AM_2_R than all other agonist-receptor pairings due to slow off-rate binding kinetics. We mapped the regions responsible for the slow off-rate to the AM2/IMD mid-region and to the RAMP3 ECD. Molecular dynamics (MD) simulations enabled by rebuilding the AM_2_R cryo-EM structures explained the structural basis for the distinct kinetics. Our results indicate that AM2/IMD-AM_2_R is a cognate pair and show how the peptide agonist and RAMP accessory protein collaborate to shape CLR signaling. We conclude that AM2/IMD is the endogenous agonist of the AM_2_R and suggest that its long-acting signaling at this receptor is the distinguishing feature of AM2/IMD.

## Results

### AM2/IMD exhibits long-duration cAMP signaling at the AM_2_R (CLR-RAMP3)

We measured cAMP in real-time upon activation of each CLR-RAMP complex transiently expressed in COS-7 cells using the BRET cAMP biosensor CAMYEL^36^. COS-7 cells lack endogenous expression of CLR and RAMPs. The cells were stimulated at room temperature with 100 nM CGRP, AM, or AM2/IMD for 15 min followed by challenge with high affinity CGRP or AM variant antagonist peptides (10 μM) that we recently developed^32^. The signal decay after antagonist addition provided a measure of signal duration and we reasoned that it would act as a proxy for agonist dissociation. Each agonist gave a rise and fall to steady-state curve at each receptor in the absence of antagonist, and the BRET signal levels reflected the expected agonist potency rank orders (Fig. 1A-C). Upon antagonist challenge, the cAMP BRET signal decayed quickly to baseline for each agonist at the CLR-RAMP1 and −2 complexes and for CGRP and AM at the CLR-RAMP3 complex (Fig. 1A-C). In contrast, the cAMP signal for AM2/IMD at the CLR-RAMP3 complex decayed substantially slower (Fig. 1C). The decay phase of each curve was best fit by a one-phase exponential decay model (Fig. S1A and B). The decay rates, halflives, and time constants are summarized in Table S1 and the half-lives are shown in a scatter plot (Fig. 1D). The half-lives were within 0.6-3.5 min, except for AM2/IMD at AM_2_R, which exhibited a half-life of ~ 17 min (Fig. 1D). A slower decay rate for AM2/IMD-AM_2_R was also observed at the physiological temperature of 37°C (Fig. S1C).

**Fig. 1.**
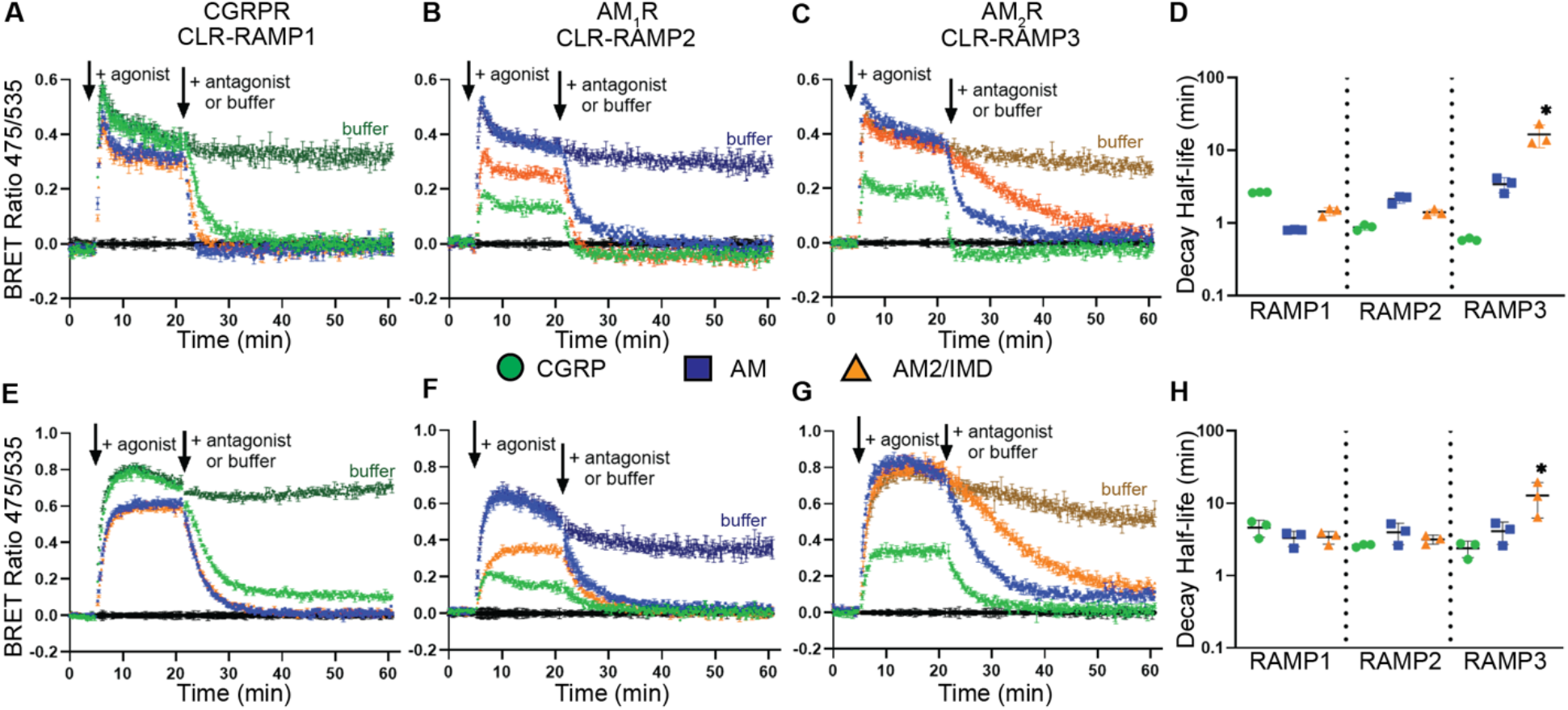
CGRP, AM, and AM2/IMD cAMP signaling kinetics. **(A-D)** COS-7 and **(E-H)** HEK293 cells expressing the indicated receptor and the cAMP biosensor were stimulated with 100 nM of CGRP (green circle), AM (blue square), or AM2/IMD (orange triangle) followed by 10 μM antagonist challenge. **(A, E)** CLR-RAMP1 with CGRP(8-37) [N31D/S34P/K35W/A36S] antagonist. CGRP with buffer addition is shown in dark green. **(B, F)** CLR-RAMP2 with AM(22-52) [S48G/Q50W] antagonist. AM with buffer addition is shown as dark blue. **(C, G)** CLR-RAMP3 with AM(22-52) [S48G/Q50W] antagonist. AM2/IMD with buffer addition is shown in brown. Plots show a representative of three independent experiments each conducted with duplicate technical replicates. Error bars show standard deviation of technical replicates. **(D, H)** Scatter plots summarizing the decay half-life for each receptor-peptide combination as mean ± SEM from three independent replicates. Star indicates significance as compared to all other combinations determined by one-way ANOVA with Tukey’s post hoc test. See Table S1 for summary of values.

We extended these experiments to HEK293 cells. Our HEK293 cells exhibited a small endogenous response to CGRP in the absence of transfected receptors, but this was not large enough to confound the results with transfected receptors (Fig. S1D). No endogenous response was observed upon AM or AM2/IMD stimulation. The cAMP signaling kinetic profiles in HEK293 cells were similar to those in COS-7 cells (Fig. 1E-H). Importantly, the slower decay of the AM2/IMD signal at AM_2_R was reproduced in the second cell line (t_1/2_ ~ 13 min) (Table S1). These results indicated that the AM2/IMD-AM_2_R pairing is unique in exhibiting a long-duration cAMP signaling capability.

### AM2/IMD is a slow off-rate, long residence time ligand of the AM_2_R

To test if AM2/IMD binding kinetics were responsible for its long-acting signaling, we used nanoBRET technology^37^ to compare the binding of AM and AM2/IMD to the AM_2_R. CLR was tagged with the BRET donor nanoluciferase (Nluc) at its N-terminus and we designed and ordered synthetic AM and AM2/IMD peptides labeled with the acceptor fluorophore TAMRA on a Lys residue substituted at equivalent positions in AM (N40) and AM2/IMD (D35) (Fig. 2A). These residues were chosen based on their solvent-exposed locations in the crystal and cryo-EM structures^30,31,34^ and mutagenesis studies, which indicated that their substitution did not alter receptor ECD binding affinity^31,38^. Wild-type pharmacology was observed for the Nluc-CLR-RAMP3 receptor with wild-type peptides (Fig. S2A) and for the TAMRA-labeled peptides at untagged CLR-RAMP3 (Fig. S2B) in cAMP accumulation assays. We further tested the labeled peptides in the real-time cAMP assay, which revealed wild-type behavior for AM-TAMRA (t_1/2_ ~ 3 min) and a gain-of-function (slower decay) for AM2/IMD-TAMRA (t_1/2_ ~ 43 min) (Fig. S2C,D). These experiments indicated that the tagged receptor and peptides exhibited wild-type or near wild-type pharmacology and were thus suitable for nanoBRET studies.

**Fig. 2.**
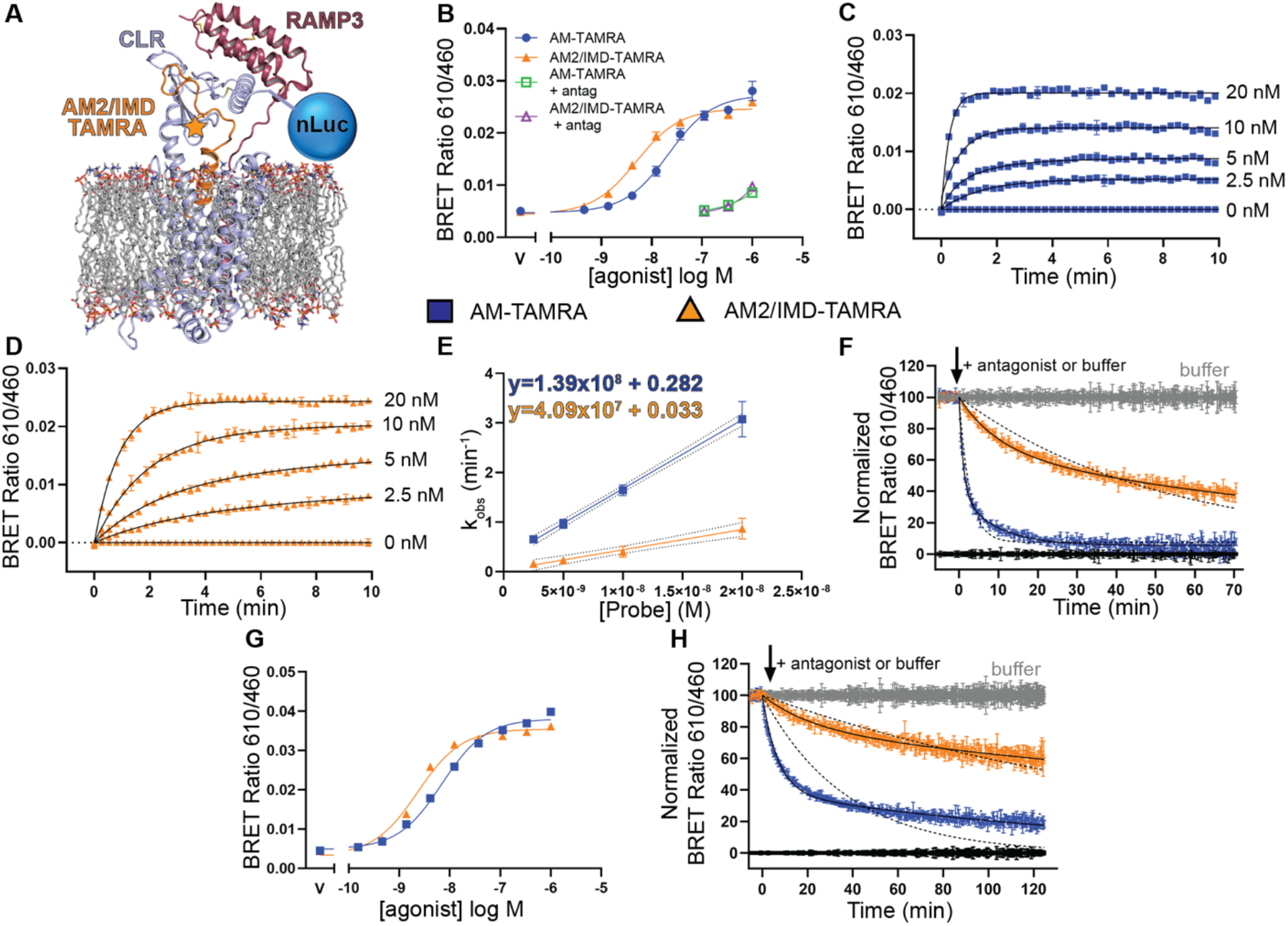
nanoBRET binding kinetics for AM-TAMRA and AM2/IMD-TAMRA with NLuc-CLR:RAMP3 membranes. **(A)** Cartoon depicting the positions of the NLuc donor and TAMRA acceptor (star) labels. **(B)** Equilibrium binding for the indicated peptides in the absence or presence of 10 μM AM(22-52) [S48G/Q50] antagonist in the presence of 50 μM GTPγS. **(C, D)** Association kinetics of AM-TAMRA (C) and AM2/IMD-TAMRA (D) in the presence of 50 μM GTPγS. **(E)** Observed rate vs. probe concentration plot for the indicated probes. **(F)** Dissociation kinetics for 10 nM AM-TAMRA or AM2/IMD-TAMRA with dissociation initiated by 1 μM AM(22-52) [S48G/Q50W] in the presence of 50 μM GTPγS. Curves were fit to a two-phase exponential decay (solid line) and a one-phase exponential decay (dashed lines). **(G)** As in (B) except in the presence of 30 μM miniGs. **(H)** As in (F) except in the presence of 30 μM miniGs. All plots show a representative of three independent experiments with duplicate technical replicates. Error bars show standard deviation for technical replicates.

Nluc-CLR was co-expressed with RAMP3 in COS-7 cells and membranes were prepared for the binding studies, which were conducted at 25°C and in the presence of GTPγS to uncouple the receptor from G protein. Equilibrium binding experiments revealed saturable binding of AM-TAMRA and AM2/IMD-TAMRA with binding affinities of 26 and 7 nM, respectively (Fig. 2B, Table S2). Varying the incubation time indicated that 3 hr was sufficient to reach equilibrium (Fig. S2E). In association kinetics experiments AM-TAMRA reached equilibrium quicker than AM2/IMD-TAMRA (Fig. 2C, D). Extended incubation revealed signal decay that we were unable to eliminate or correct for (Fig. S2F, G), so we limited analysis of the association data to the first 10 min. The individual curves were best fit by a one-phase association exponential model and plots of the observed rates vs. probe concentration showed a linear relationship consistent with a single-step binding model (Fig. 2E). The on- and off-rates were determined as the slope and y-intercept, respectively, as described^39^. This revealed slower on- and off-rates and a longer half-life for AM2/IMD-TAMRA (t_1/2_ ~ 21 min) as compared to AM-TAMRA (t_1/2_ ~ 2.5 min) (Table S2). The Kd values calculated from the on-and off-rates were ~ 10-fold lower than the equilibrium Kd values. These discrepancies may reflect inaccuracies in fitting the kinetic data due to the signal decay issue and/or indicate that a single-step binding model does not appropriately describe these interactions.

Next, we turned to dissociation kinetics experiments. The membranes were incubated with 10 nM of each probe for 25 min to reach equilibrium followed by injection of either buffer or 1 μM high-affinity unlabeled AM variant antagonist to initiate dissociation. Signal decay was also evident in these experiments (Fig. S2H), but the decay could be corrected for by normalization to the buffer control injection. These data revealed much slower dissociation of AM2/IMD-TAMRA than AM-TAMRA and the dissociation curves were best fit by a two-phase exponential decay model (Fig. 2F). AM-TAMRA exhibited fast and slow component half-lives of 0.75 and 6.9 min, respectively, whereas AM2/IMD-TAMRA had fast and slow component half-lives of 7 and 76 min, respectively (Table S2). The fast components for AM-TAMRA and AM2/IMD-TAMRA accounted for 67% and 35% of the dissociation curves, respectively. These data are consistent with a two-step (un)binding process as proposed for other class B GPCR peptide ligands^40–42^, and indicate that AM2/IMD-TAMRA has a substantially longer residence time than AM-TAMRA at the uncoupled AM_2_R.

We also performed equilibrium binding and dissociation kinetics experiments in the presence of purified sumo-mini-Gs fusion protein, which is a G protein surrogate that locks the receptor in the active, coupled state^17^. The two probes exhibited higher affinity in the presence of mini-Gs, but the fold increase was slightly larger for AM-TAMRA than AM2/IMD-TAMRA, which had affinities of 8 and 3 nM, respectively (Fig. 2G, Table S2). Both probes yielded biphasic dissociation curves for the G protein-coupled state of AM_2_R and AM2/IMD-TAMRA again exhibited substantially slower dissociation than AM-TAMRA (Fig. 2H, Table S2).

### AM2/IMD promotes more stable CLR-RAMP3 complexes than AM

We previously reported a membrane protein native PAGE mobility shift assay for the biochemical characterization of agonist-dependent coupling of CLR-RAMP1-3 complexes to mini-Gs^17,43^. Membranes co-expressing EGFP-tagged CLR and a RAMP are incubated with agonist and purified mini-Gs, solubilized with LMNG/CHS, and analyzed by native PAGE. The heterodimeric CLR-RAMP and quaternary agonist-CLR-RAMP-miniGs complexes exhibit different mobilities due to their different sizes (Fig. 3A). AM and AM2/IMD promoted formation of CLR-RAMP3-mini-Gs complexes with equal potencies in this assay^17^. Here, we compared the stabilities of the two quaternary complexes to disruption by antagonist at 4°C. This allowed us to compare the unlabeled wild-type agonists. The membranes were incubated with AM or AM2/IMD (200 nM) and excess mini-Gs for 30 min followed by solubilization to form the quaternary complexes. The complexes were then challenged with a high-affinity AM variant antagonist for 2 or 19 hrs followed by native PAGE. The antagonist dose-dependently disrupted the AM quaternary complex with similar results observed with 2 or 19 hr antagonist exposure (Fig. 3B). Some breakdown of the AM complex was evident even in the absence of antagonist with the 19 hr incubation, indicating complex instability with extended incubation time. In contrast, the AM2/IMD quaternary complex was more resistant to breakdown in the presence of the antagonist (Fig. 3C). Remarkably, even after 19 hr exposure to 10 μM antagonist there was some AM2/IMD complex remaining. These results are consistent with the nanoBRET experiments for the coupled state of AM_2_R.

**Fig. 3.**
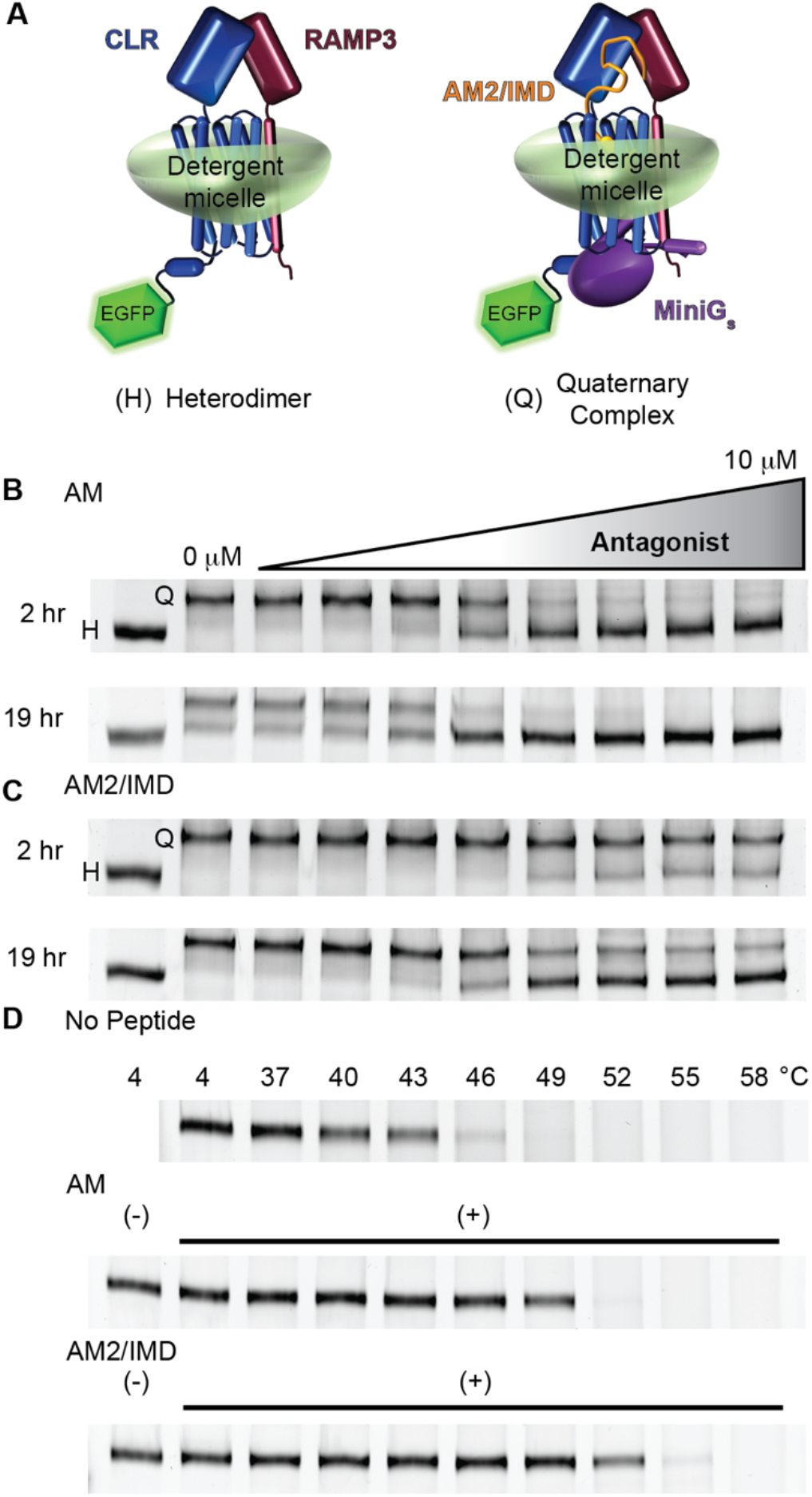
Stability of CLR-RAMP3 complexes analyzed by native PAGE. **(A)** Cartoon depicting detergent-solubilized heterodimer and quaternary complexes. **(B, C)** Membranes expressing MBP-CLR-EGFP and MBP-RAMP3 were incubated with 200 nM agonist and 50 μM miniGs to form solubilized quaternary complexes followed by challenge with 3-fold serial dilutions of AM(22-52) [S48G/Q50W] antagonist for the indicated times. **(D)** Heterodimer thermostability assay. The membranes were incubated at the indicated temperatures in the absence or presence of the indicated peptides (1 μM) followed by native PAGE analysis. In all panels, the heterodimer and quaternary complexes were resolved on 8% hrCNE native gels and imaged for in-gel EGFP fluorescence. Gels shown are a representative of three independent experiments.

Next, we tested the thermostability of CLR-RAMP3 when bound to AM or AM2/IMD in the absence of mini-Gs. The membranes were incubated in the absence or presence of 1 μM AM or AM2/IMD for 30 min followed by solubilization. The solubilized complexes were then incubated an additional 30 min at various temperatures followed by centrifugation and analysis of the supernatants by native PAGE. The agonists alone are too small to induce a mobility shift, but their binding was evidenced by the increased thermostability of the CLR-RAMP3 complex in their presence (Fig. 3D). The ligand-free AM_2_R was stable up to ~43°C, whereas the AM- and AM2/IMD-bound AM_2_R were stable to ~49°C and ~52°C, respectively. The greater stability in the presence of AM2/IMD is consistent with it having a slower off-rate at the uncoupled AM_2_R as observed in the nanoBRET experiments.

### The AM2/IMD mid-region is responsible for its long-acting behavior at AM_2_R

To determine the region of AM2/IMD responsible for its slow off-rate at AM_2_R, we examined chimeras of AM and AM2/IMD in the real-time cAMP signaling assay in COS-7 cells. First, we tested two previously described chimeras^17^ with the junction at the half-way point at the end of the N-terminal α-helix (Fig. 4A). We refer to these as AM-AM2 half and AM2-AM half (Fig. 4B). These exhibited decay half-lives similar to AM and AM2/IMD at the CGRPR and AM_1_R, and similar to AM at the AM_2_R (Fig. 4C-E, N-P; Table S3). Their lack of slow decay at the AM_2_R indicated that the N- and C-terminal halves of AM2/IMD were insufficient to confer the slow decay rate. Next, we tested chimeras with a junction at the end of the central “hinge” region that connects the TMD and ECD binding portions (Fig. 4A). We refer to these as AM-AM2 ECD and AM2-AM ECD (Fig. 4F). AM-AM2 ECD exhibited decay half-lives similar to AM at each receptor, whereas AM2-AM ECD behaved like AM2/IMD at the CGRPR and exhibited a gain-of-function slow decay at both AM_1_R (t_1/2_ ~ 6 min) and AM_2_R (t_1/2_ ~ 33 min) (Fig. 4G-I, N-P; Table S3). These results suggested that the slow decay required the AM2/IMD hinge region and additional residues prior to the hinge that were lacking in AM-AM2 half.

**Fig. 4.**
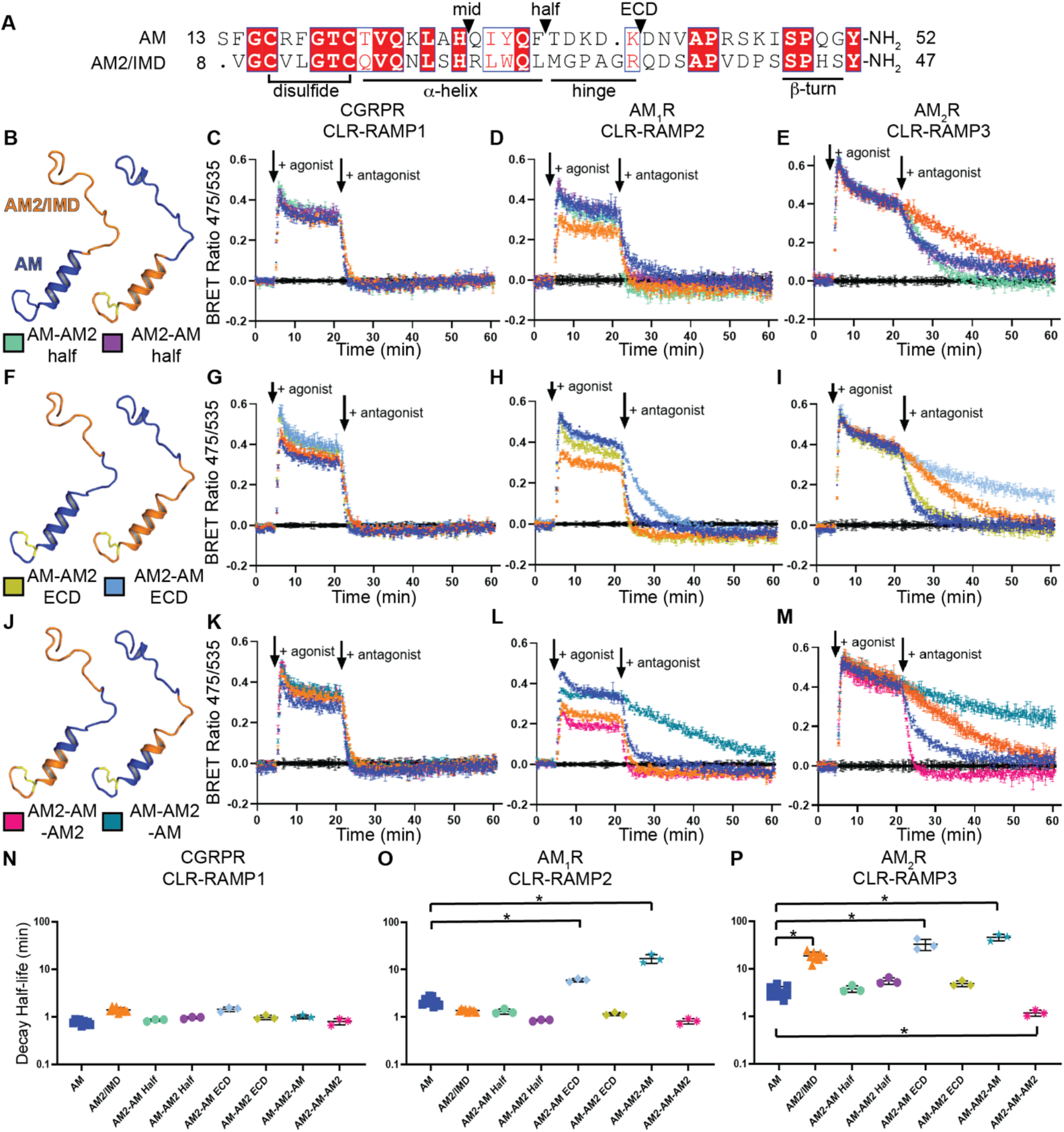
Mapping the AM2/IMD region responsible for slow cAMP signaling decay. **(A)** Amino acid sequence alignment of AM(13-52) and AM2/IMD(8-47). Triangles indicate chimera junction points. **(B)** Structural depiction of AM-AM2 half and AM2-AM half chimeras tested in C-E. **(C-E)** cAMP signaling kinetics for the indicated receptors in COS-7 cells stimulated with 100 nM of the indicated agonist followed by 10 μM antagonist. For CLR-RAMP1, CGRP(8-37) [N31D/S34P/K35W/A36S] antagonist was used and AM(22-52) [S48G/Q50W] antagonist was used at CLR-RAMP2/3. **(F)** Structural depiction of AM-AM2 ECD and AM2-AM ECD chimeras tested in G-I. **(G-I)** As in C-E with the ECD chimeras. **(J)** Structural depiction of AM2-AM-AM2 and AM-AM2-AM chimeras tested in K-M. **(K-M)** As in C-E with the mid-region chimeras. In all plots, wild-type AM and AM2/IMD control agonists are blue and orange, respectively, and the chimeras are colored according to the legend in B, F, and J. All plots are shown as mean ± SD for technical replicates for a single representative of three independent experiments. **(N-P)** Scatter plots summarizing the decay half-lives of all the chimeric peptides compared to WT AM and AM2/IMD. Statistical significance was determined by one-way Anova with Tukey’s Post Hoc test. See Table S3 for the summary of the decay kinetic values.

To define the central AM2/IMD segment responsible for the slow decay, we tested chimeras with swapped mid-regions. The N-terminal junction was after a conserved His in the α-helix (“mid” in Fig. 4A) and the C-terminal junction was at the end of the hinge. The “mid” junction point was chosen because AM2/IMD R23 forms a salt bridge with D96 at the “bottom” of the CLR ECD^34^ and we hypothesized that this interaction contributes to the slow decay. We refer to these chimeras as AM2-AM-AM2 and AM-AM2-AM (Fig. 4J). AM2-AM-AM2 exhibited decay similar to AM at the CGRPR, and loss-of-function faster decay (as compared to AM) at both AM receptors (Fig. 4K-P; Table S3). AM-AM2-AM had fast decay at the CGRPR and striking gain-of-function slower decay as compared to AM at AM_1_R (t_1/2_ ~ 17 min) and as compared to AM2/IMD at AM_2_R (t_1/2_ ~ 49 min) (Fig. 4K-P; Table S3). These results indicated that the AM2/IMD central 11 residue segment from R23 to R33 is responsible for its slow off-rate at AM_2_R.

### The RAMP3 ECD enables the long-acting behavior of AM2/IMD

To identify the RAMP3 region that enables the AM2/IMD slow off-rate, we constructed six chimeras of RAMP2 and RAMP3 and examined their properties in the real-time cAMP signaling assay in COS-7 cells. We swapped the ECD, transmembrane helix (TMD), or C-terminal cytoplasmic tail using the junction points in Fig. 5A. The chimeras were co-expressed with CLR and the AM and AM2/IMD signaling kinetics were assessed. Strikingly, swapping the RAMP2/3 ECD completely swapped the AM2/IMD slow decay phenotype, while having little to no effect on AM signaling kinetics (Fig. 5B-E; Table S4). In contrast, swapping the RAMP2/3 TMD (Fig. 5F-I; Table S4) or C-terminal tail (Fig. 5J-M; Table S4) had little to no effect on the signal decay for both agonists. These results indicated that the slow decay of AM2/IMD signaling at the AM_2_R is dictated by the RAMP3 ECD.

**Fig. 5.**
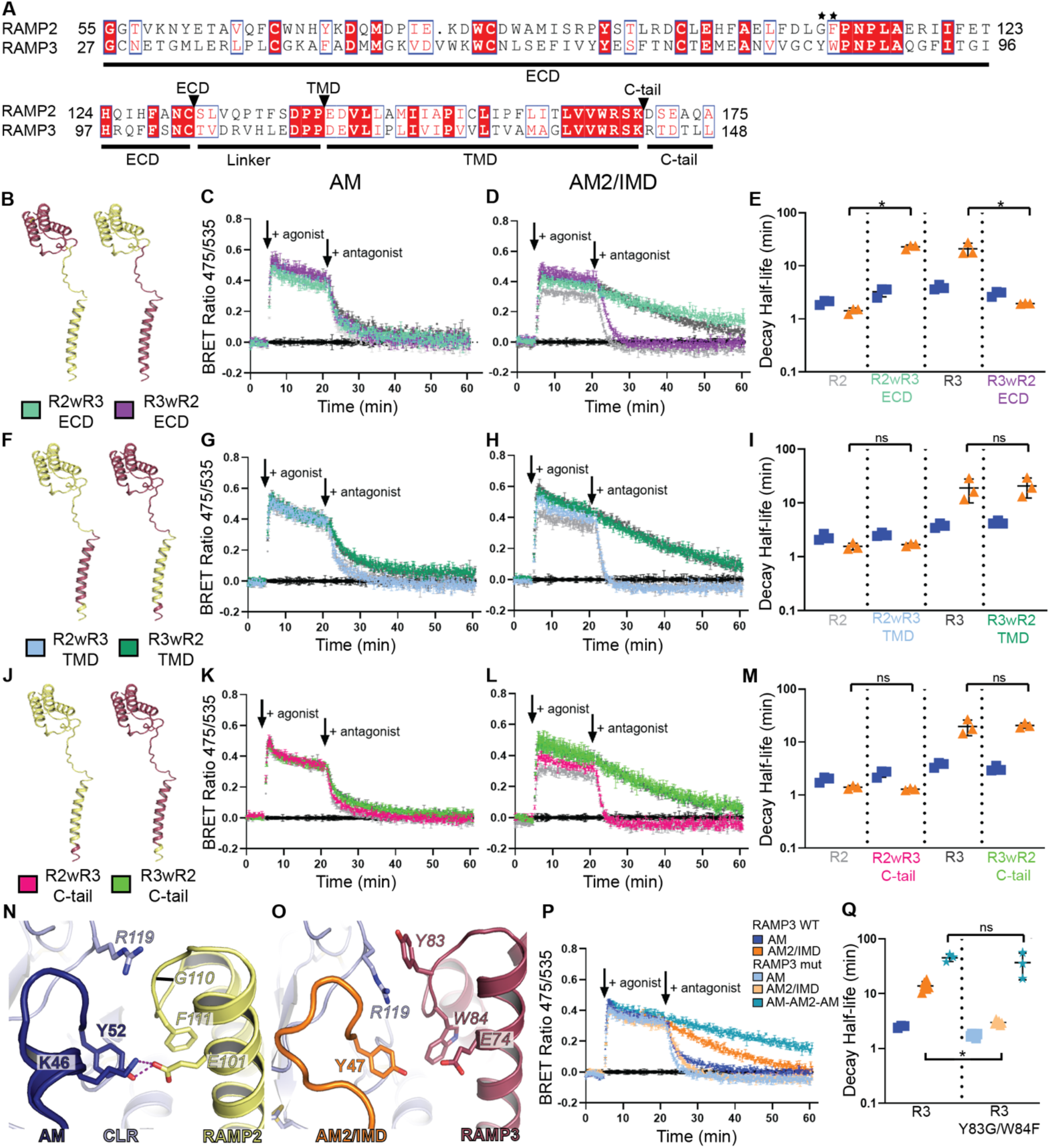
Mapping the RAMP3 region responsible for the slow AM2/IMD cAMP signaling decay. **(A)** Amino acid sequence alignment of RAMP2 and RAMP3 showing the chimera junction points. Stars indicate the residues mutated in panel P and Q. **(B)** Structural depiction of ECD swap chimeras tested in C and D with wild-type RAMP2 shown in yellow and RAMP3 shown in dark red. **(C, D)** cAMP signaling kinetics in COS-7 cells for the indicated wild-type CLR-RAMP2/3 or CLR-RAMP2/3 chimeras stimulated with 100 nM AM (C) or AM2/IMD (D) followed by 10 μM challenge with AM(22-52) [S48G/Q50W] antagonist. **(E)** Scatter plots summarizing decay half-lives for the receptors used in C and D. **(F-I)** As in B-E except with RAMP TMD swap chimeras. **(J-M)** As in B-E except with RAMP C-tail swap chimeras. In plots C-E, G-I, and K-M, wild-type RAMP2 is shown in light grey and RAMP3 in dark grey and the RAMP2/3 chimeras are colored based on the legends in panels B, F, and J. Wild-type AM is shown in blue and AM2/IMD is shown in orange. **(N)** AM bound to CLR and RAMP2 from PDB 4RWF. **(O)** AM2/IMD bound to CLR and RAMP3 from 6UVA_rebuild. **(P)** cAMP signaling kinetics for CLR-RAMP3 wild-type and Y8G/W84F stimulated with AM, AM2/IMD, or AM-AM2-AM chimera agonist followed by antagonist challenge. **(Q)** Scatter plot summarizing decay half-life for the indicated receptor-peptide combinations in P. AM-AM2-AM half-life values at WT RAMP3 were re-plotted from Fig. 4P for comparison purposes. All plots show a single representative of three independent experiments with standard deviation for duplicate technical replicates. Statistical analysis was determined with one-way ANOVA using Tukey’s Post Hoc test.

Seeking to determine the elements of the RAMP3 ECD that enabled the AM2/IMD slow off-rate, we considered distinctions among the RAMP2/3 ECD residues that augment the CLR ECD binding pocket. We previously showed that AM bound the purified RAMP2-CLR and RAMP3-CLR ECD complexes with nearly equal equilibrium affinities (K_I_ ~ 5 μM), whereas AM2/IMD bound the RAMP3-CLR ECD with an affinity (K_I_ ~ 2 μM) 7-fold stronger than the RAMP2-CLR ECD (K_I_ ~ 14 μM)^31^. These differences likely reflect the distinct RAMP2/3 residues that augment the binding site including RAMP2 G110, F111 and RAMP3 Y83, W84, which confer different contours to the binding pocket occupied by the peptide C-terminus (Fig. 5N, O). Notably, RAMP2/3 share E101/E74, which are important for binding AM^30,32,44,45^. In the RAMP2 structure E101 directly contacts AM K46 and Y52. We reasoned that a RAMP3 Y83G/W84F double swap mutant would negatively affect AM2/IMD more than AM, so we characterized this mutant. AM and AM2/IMD were equipotent at the AM_2_R with RAMP3 Y83G/W84F in a cAMP accumulation assay, albeit with slight potency reductions as compared to wild type receptor (Fig. S3). In the CAMYEL cAMP signaling assay, AM kinetics were nearly wild type at the mutant receptor, whereas AM2/IMD exhibited dramatically faster signal decay (t_1/2_ ~ 3 min) that was nearly equal to that of AM (Fig. 5P, Q; Table S4). In contrast, the AM-AM2-AM chimera, which contains the AM ECD binding region, retained its slow signal decay phenotype at the RAMP3 mutant AM_2_R (t_1/2_ ~ 37 min) (Fig. 5P, Q). These results indicated that the RAMP3 ECD enabled the AM2/IMD long-acting signaling primarily via its role in forming the binding site for the peptide C-terminus.

### MD simulations reveal the structural basis for the AM2/IMD slow off-rate

We sought to perform MD simulations to compare the AM- and AM2/IMD-bound AM_2_R structures. This required resolving the issue of the different ECD-binding conformations of AM and AM2/IMD in the cryo-EM^34^ and crystal structures^30,31^. We rebuilt and re-refined the AM (6UUS) and AM2/IMD (6UVA) AM_2_R cryo-EM structures by using AlphaFold2^46^ to obtain a RAMP3 ECD model combined with manual rebuilding as guided by fitting to the deposited cryo-EM density maps. This yielded structures with improved fits to the maps and showed that the AM and AM2/IMD ECD-binding conformations in the cryo-EM structures were the same as those observed in the prior crystal structures (Fig. S4). These improved structures served as the starting points for the MD simulations.

A set of three parallel molecular simulations were run for each of the AM-AM_2_R and AM2/IMD-AM_2_R systems. Each system included the full CLR and RAMP3 proteins and the bound peptide (Fig. 6A). The complex was embedded in an explicit membrane (40% cholesterol, 60% POPC) and included restraints on the intracellular CLR residues to mimic the presence of the G protein. To achieve better sampling of molecular conformations, we used a variant of the weighted ensemble algorithm^47^ called REVO (“Resampling Ensembles by Variation Optimization”)^48^ that periodically merges and clones trajectories in order to achieve a more diverse set of structures. The criteria we chose to measure diversity was the root mean squared distance (RMSD) of the peptide after alignment to the entire receptor dimer, which has been previously used to generate ligand unbinding trajectories^49-51^. In the simulations here, both the TM and EC regions of the peptide remained associated with the receptor throughout, although the linker regions were flexible, allowing the ECD of the receptor to move significantly with respect to the TMD in both cases (Fig. 6B).

**Fig. 6.**
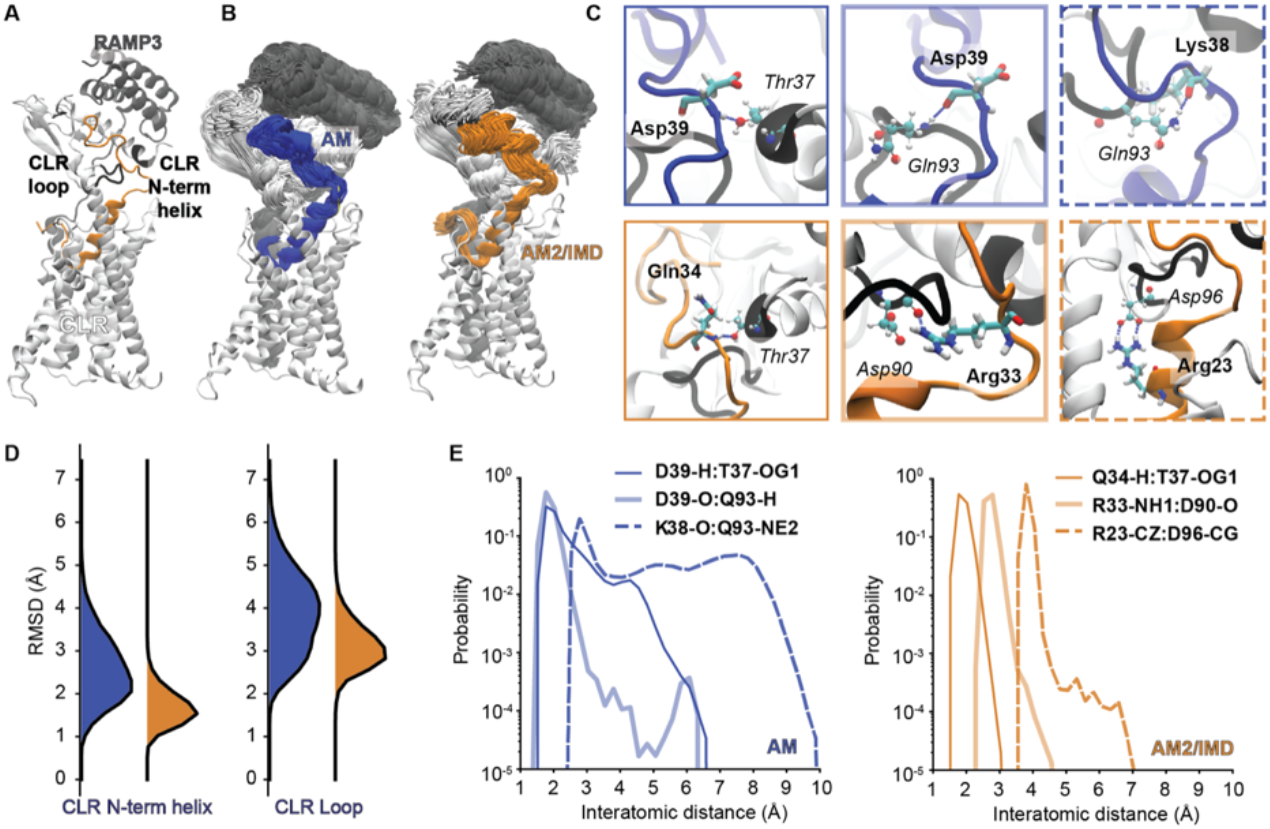
Molecular dynamics analysis of AM and AM2/IMD mid regions. **(A)** A system overview shows the CLR protein (white), RAMP3 (grey) and the AM2/IMD peptide (orange). Two regions of CLR are labeled: the loop region (residues 90-97) and the N-terminal helix (residues 36-38). These are shown in black in panel (C). **(B)** An overlay of all 144 final structures from the REVO simulations are shown for AM (blue, left) and AM2/IMD (orange, right). Structures were aligned using the CLR TMD region and only one representation of the CLR TMD is shown for clarity. **(C)** The three intermolecular hydrogen bonding interactions that were the most stable for each system are shown, with the specific residues in licorice representation and a dashed blue line showing the hydrogen bonds. Peptide residues are labels in bold and CLR residues are labeled in italics. The linetype of the borders correspond to the probability curves in panel (E). **(D)** Probability distributions of the root mean squared distance (RMSD) of specific CLR regions after alignment to the peptide mid-region for both AM (blue, left) and AM2/IMD (orange, right). **(E)** Probability distributions of distances reporting on the most stable hydrogen bonding interactions in each system for AM (blue, left) and AM2/IMD (orange, right). The specific atoms used for the distance calculation were chosen to be optimal in the case of symmetry and are reported in the legend.

Despite the significant motion of the ECD with respect to the TMD, the hinge region of the peptide maintained stable hydrogen bonding interactions with residues in the CLR ECD loop (residues 90 to 97) and the CLR N-terminal helix (residues 36 to 38), both shown in black in Fig. 6A. The most stable interactions for each peptide are analyzed in Fig. 6C and 6E. AM2/IMD maintains these interactions with higher probability. R33 played a central role in coordinating interactions with D90 of the CLR ECD loop, as well as intramolecular hydrogen bonds with the backbone atoms of P30 and M28. R23 formed a strong salt-bridge interaction with D96 at the bottom of the CLR ECD; no analogous residue is present in AM. These interactions in AM2/IMD-AM_2_R work together to stabilize the hinge region. After alignment to the hinge region, AM2/IMD-AM_2_R showed lower RMSDs of both the CLR N-terminal helix and the CLR ECD loop than AM-AM_2_R (Fig. 6D). This reveals a more stable bound complex which could contribute to the longer observed residence time of AM2/IMD.

Fig. 7 analyzes the flexibility and molecular interactions in the ECD complexes. By aligning the set of 144 final structures from the REVO-MD simulations using the CLR ECD, we observed a slight increase in stability of the C-terminal end of AM2/IMD in comparison to AM (Fig. 7A). A quantitative analysis is provided in Fig. 7B, where we analyze the entire ensemble of AM and AM2/IMD conformations using different RMSD measurements and alignments. For each domain, with each possible alignment, we found larger conformational changes in the AM-bound system in comparison to the AM2/IMD-bound system. To further investigate the stabilizing interactions observed in the MD simulations, we show representative high-weight frames from the end of the REVO-MD simulation (Fig. 7C). We also computed the density of water molecules after alignment to a set of residues surrounding the terminal tyrosine in the peptide and W84 of RAMP3. The highest probability water-binding sites in the region are shown by a transparent red surface. The AM2/IMD-bound system was stabilized by a set of water-mediated hydrogen bonds involving G71 (CLR), Y47 (AM2/IMD), E74 (RAMP3) and the backbone nitrogen of W84 (RAMP3). This involved three labeled water molecules, which all reside in the highest-density solvation pockets. Note that K46 of AM, while it introduces a powerful salt-bridge interaction, disrupts this water-mediated H-bond network. Significantly, we also observed a stable hydrogen bond between the side chains of W84 and T73. While this is present in both systems, it is more stable in the AM2-IMD system (Fig. 7D), likely due to the stabilizing interactions afforded by the water-mediated hydrogen bond network. This is consistent with the effects of the RAMP3 W84F mutation described above, which would eliminate this H-bond and destabilize the water-mediated H-bond network that is specific to the AM2/IMD system.

**Fig. 7.**
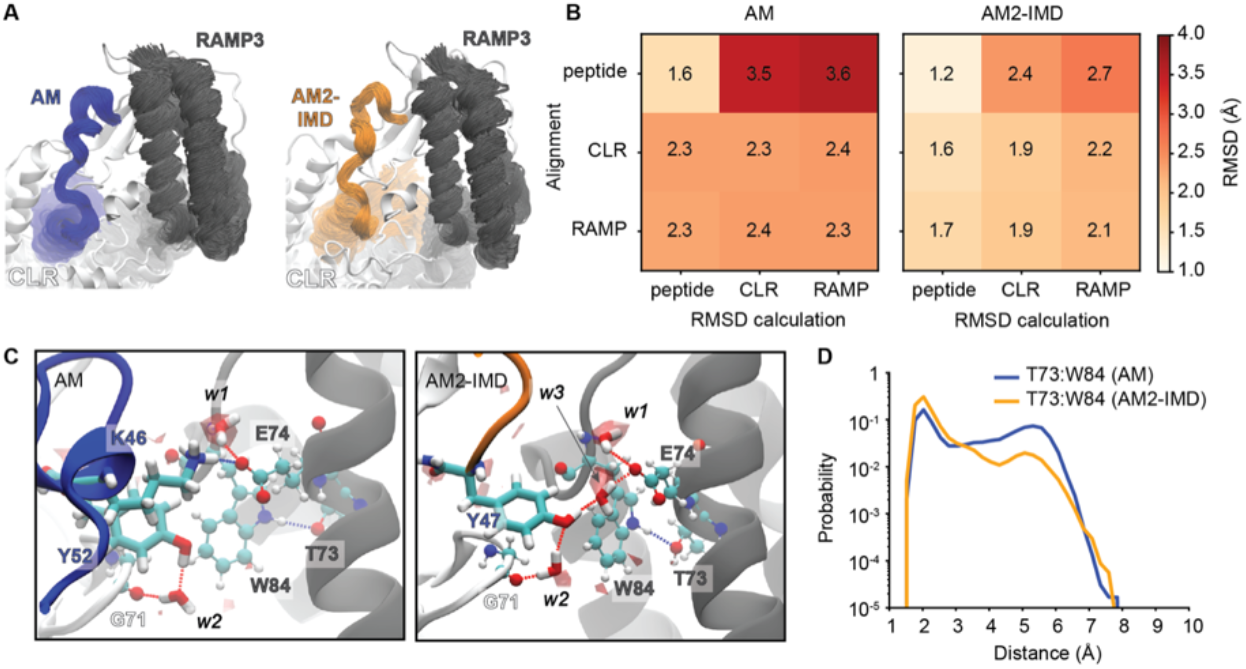
Molecular dynamics analysis of AM and AM2/IMD extracellular regions. **(A)** A visualization of the conformational heterogeneity of the extracellular domains is shown for AM (left) and AM2-IMD (right). In both cases, structures are aligned using CLR residues 35-123. **(B)** Average RMSD calculations of: the peptide, the CLR ECD (residues 35-123) and the RAMP3 ECD (residues 29-102) are shown after three different alignments. Averaged RMSDs are calculated after alignment to each of the same three regions and the results are shown in a 3×3 matrix. The reference structure used in each case is the structural model used to initialize the dynamics. **(C)** The peptide C-terminal binding site for AM (left) and AM2-IMD (right). Peptide residues and water molecules are shown in licorice and residues of CLR and RAMP are shown in ball and stick representation. Water density isosurfaces are shown using transparent red. The isosurfaces were computed using the volmap tool of VMD and the same isovalue of 0.903 was used to visualize each density. **(D)** Probability distributions for the RAMP3 T73:W84 intramolecular hydrogen bond shown in panel C.

## Discussion

For nearly two decades AM2/IMD has been thought to be a relatively non-selective agonist of the three CLR-RAMP complexes^1,19,27^. Consistent with this, AM2/IMD has many actions in common with CGRP and AM, so the rationale for the existence of a third agonist for the CGRP and AM receptors has been unclear. Similarly, the CGRP and AM1 receptors have well defined roles in mediating key functions of CGRP and AM, respectively, but the reason for a second AM receptor, AM_2_R, and its functions are poorly understood. Here, we discovered that the AM2/IMD-AM_2_R pairing has kinetic features that set it well apart from the eight other agonist-receptor combinations. Rather than being non-selective, AM2/IMD is kinetically selective for the AM_2_R at which it elicits long-duration cAMP signaling. We propose that AM2/IMD is the endogenous agonist of the fortuitously named AM_2_R and suggest that the primary role of the AM_2_R is to mediate long-duration AM2/IMD signaling.

Our discovery was facilitated by the cAMP biosensor assay in combination with the engineered high affinity peptide antagonists that we recently developed^32^ (Fig. 1). This facilitated straightforward characterization of the signaling duration capability of any agonist without the need for wash-out steps and allowed rigorous quantitation of the signal decay rates, which we reasoned would be a proxy for agonist off-rate. The nanoBRET binding assays bore this out by showing that AM2/IMD-TAMRA dissociated slowly from both the uncoupled and coupled states of AM_2_R (Fig. 2, Table S2). NanoBRET technology is powerful, but its limitations were evident in the association kinetics experiments where the calculated Kd values were ~10-fold lower than the equilibrium affinities. These discrepancies may reflect the signal decay issue, insufficient time resolution to define the early portion of the curves, and/or insufficient probe concentration range due to the decay being more pronounced at high probe concentrations. These issues may have prevented detection of the biphasic association curves expected for a 2-step (un)binding mechanism^41^. PTH and PTHrP exhibited biphasic association curves for the class B parathyroid hormone receptor (PTH1R) using a different assay technology^40,42^. Similar nanoBRET association kinetics discrepancies with equilibrium binding were reported in a study of relaxin binding to its GPCR, which involves a multi-step mechanism^52^.

Fortunately, in the nanoBRET dissociation kinetics experiments we could correct for signal decay, which revealed two-phase AM-TAMRA and AM2/IMD-TAMRA dissociation curves. For the uncoupled AM_2_R, the fast unbinding component may arise from peptide-receptor complexes in which the peptide is engaged solely to the receptor ECD and the slow unbinding component may come from complexes in which the peptide is fully engaged to both ECD and TMD. This interpretation is consistent with a cryo-EM structure of the CGRP-bound CGRP receptor in the absence of G protein^53^. This showed that CGRP was primarily engaged to the receptor ECD, with a fraction of complexes showing limited engagement of the TMD by CGRP. Here, the fast dissociation components for AM-TAMRA and AM2/IMD-TAMRA accounted for 66.5% and 34.7% of their curves, respectively, which implies that AM2/IMD-TAMRA had a greater capacity to fully engage the uncoupled AM_2_R than AM-TAMRA. This may explain the greater thermostability of AM2/IMD-AM_2_R observed in the native PAGE assay (Fig. 3D). AM2/IMD also had a slower off-rate from the G protein-coupled state of AM_2_R (Fig. 2H and 3C). Overall, the nanoBRET and native PAGE assays were consistent with the longer residence time of AM2/IMD at AM_2_R being responsible for its longer duration cAMP signaling capability.

Using peptide chimeras, we mapped the region responsible for the slow off-rate to the 11 amino acid R23-R33 segment of AM2/IMD that binds at the interface of the CLR ECD and TMD (Fig. 4 and 6). The MD simulations revealed a series of polar interactions stabilizing this segment and its interactions with CLR. These are anchored at one end by the intermolecular R23-CLR D96 salt-bridge and at the other by a series of H-bonds involving R33 including intermolecular bonds to the loop at the base of the CLR ECD, and intramolecular bonds to the backbones of P30 in the hinge and M28 at the end of the helix. Neither of these anchoring interactions alone was sufficient to confer the slow off-rate as revealed by the AM-AM2 half and AM2-AM half chimeras. The corresponding central segment of AM exhibited much less stability in the MD simulations. AM lacks an R23 equivalent so it cannot form the salt-bridge and its R33 equivalent is K38, which cannot form the same pattern of H-bonds due its shorter side chain and the AM hinge being one amino acid shorter than the AM2/IMD hinge. A recent study of the amylin receptors, which are RAMP complexes with the calcitonin receptor, suggested that a central 7 amino acid segment of amylin contributed to amylin receptor selectivity^54^. This study and our findings here make it clear that the mid-regions of the CGRP family peptides are not simply passive linkers.

Remarkably, the AM2-AM ECD and AM-AM2-AM chimeras both exhibited a gain-of-function slow signaling decay phenotype at the AM_1_R. Yet none of the peptides exhibited slow signaling decay capacity at the CGRPR. Why is this? This can be understood by considering the orientations of the CLR ECD relative to the TMD that are induced by each RAMP (Fig. S5A, B). RAMP2 and −3 promote similar CLR ECD-TMD arrangements^34^ such that the R23-R33 segment could form interactions in AM_1_R like those observed in AM_2_R. In contrast, RAMP1 induces a different ECD-TMD arrangement^35,53^ that would prevent the R23-R33 segment from making the same pattern of interactions (Fig. S5C-F). These different CLR ECD-TMD arrangements were attributed to the different RAMP linker sequences that connect the ECD and TM helix and were proposed to contribute to ligand selectivity^34^. Our data are consistent with this for the CGRP receptor vs. the two AM receptors, but the RAMP linkers do not control peptide selectivity among the two AM receptors (see below).

Why is AM2/IMD kinetically selective for AM_2_R if its R23-R33 segment has the capacity for slow decay at AM_1_R? This was explored through RAMP2/3 chimeras, RAMP3 mutants, and the MD simulations (Fig. 5 and 7). The chimeras showed that the RAMP3 ECD enabled the AM2/IMD slow signaling decay, and the RAMP3 Y83G/W84F swap mutant indicated that this was due to its role in forming the binding pocket for the peptide C-terminus. The simulations revealed a more stable AM2/IMD-AM_2_R ECD complex as compared to the AM-bound version, particularly for the peptide C-terminal region near RAMP3 Y83 and W84. A network of water-mediated H-bonds in the pocket involved AM2/IMD Y47 and RAMP3 E74 and is stabilized by hydrogen bonds with the RAMP3 W84 backbone as well as an intramolecular RAMP3 W84-T73 hydrogen bond. In contrast, AM binding was more dependent on the AM K46-RAMP3 E74 salt-bridge, consistent with prior mutagenesis studies^44,45^. RAMP2 can also form this salt-bridge via E101^30^, but G110 and F111 are unable to provide the optimal environment for the AM2/IMD C-terminus. This explains why AM has nearly equal affinities for the purified AM_1_R and AM_2_R ECD complexes, whereas AM2/IMD has higher affinity for the AM_2_R ECD complex^31^. Hence, the slow decay phenotype of AM2/IMD was lost at the AM_2_R with RAMP3 Y83G/W84F, whereas the AM-AM2-AM chimera retained slow decay at the mutant AM_2_R. These results highlight the substantial contribution of the CLR-RAMP ECD complexes to ligand selectivity, which makes sense because ECD-binding is likely the first step in the binding mechanism and the purified CLR-RAMP1/2/3 ECD complexes exhibited peptide selectivity profiles similar to the intact receptors^30,31,38^.

Kinetic selectivity of AM2/IMD for AM_2_R is a two-step process beginning with its preference for the AM_2_R ECD over the AM_1_R ECD, after which its mid-region R23-R33 segment enables the stable interactions at the ECD-TMD interface (Fig. 8). Together, these strong ECD and ECD-TMD interface interactions yield the slow off-rate. The weaker AM_1_R ECD binding of AM2/IMD prevents a slow off-rate at the AM_1_R. AM2/IMD binds the CGRPR ECD with the same affinity as the AM_2_R ECD^31^, but its mid-region cannot form stable ECD-TMD interface interactions due to the altered ECD-TMD arrangement in the CGRPR. Our findings emphasize the concept that the peptide agonists and the RAMP accessory proteins can collaborate to dictate CLR pharmacology. The RAMPs play a substantial role in determining receptor phenotype through a combination of their augmentation of the CLR ECD binding site and their modulation of CLR ECD-TMD inter-domain arrangement and/or dynamics, but unique agonist properties can further shape the signaling outcomes as shown here for AM2/IMD.

**Fig. 8.**
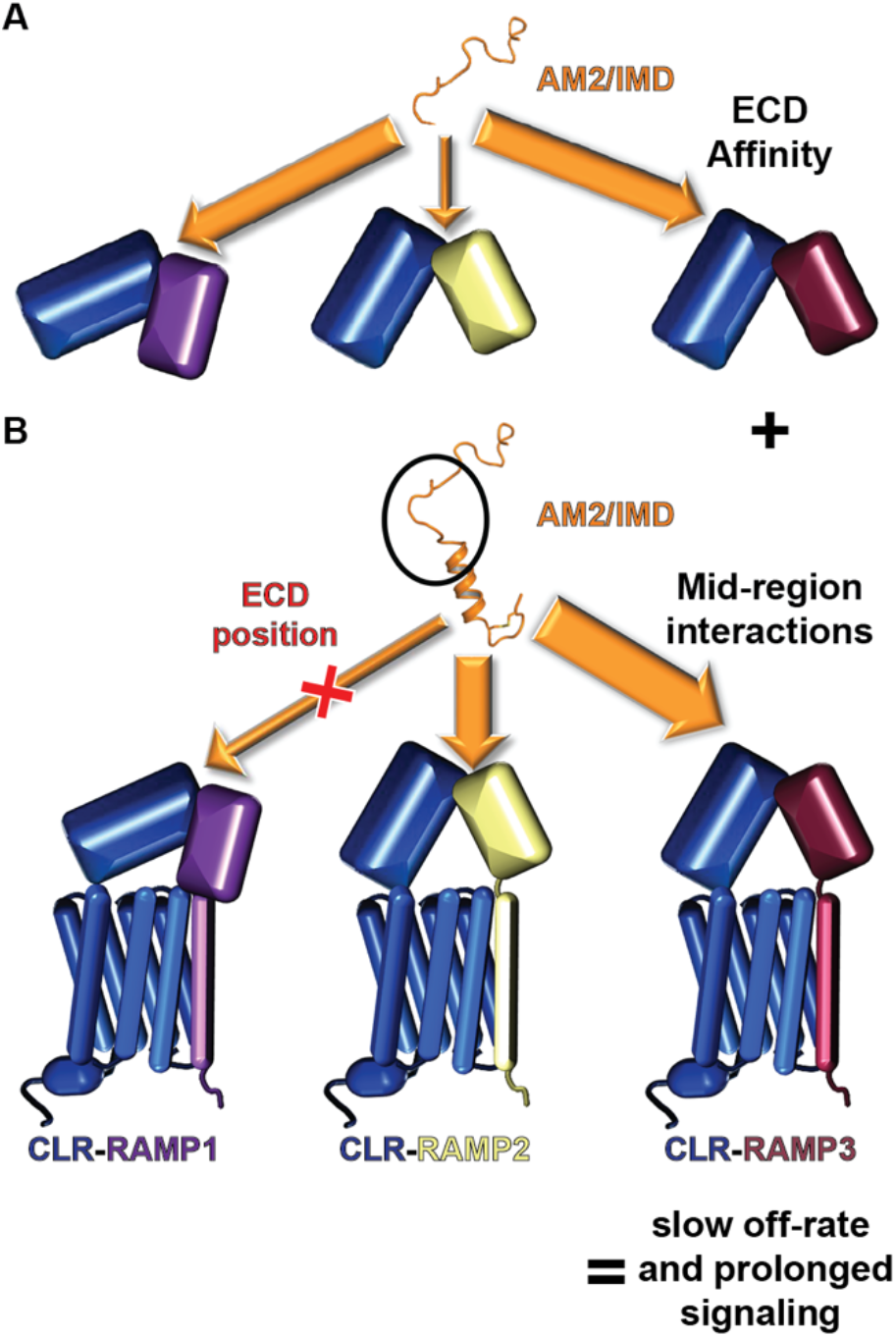
Mechanism of AM2/IMD kinetic selectivity for AM_2_R. **(A)** Cartoon depicting the strength of the interactions of AM2/IMD with the three RAMP-CLR ECD complexes. **(B)** Cartoon depicting the interactions of the AM2/IMD mid-region with each of the three RAMP-CLR complexes. The combination of strong ECD affinity and strong midregion interactions yields a slow off-rate and long duration signaling.

The pharmacology of AM2/IMD and AM at the AM_2_R is reminiscent of PTH and PTHrP at the PTH1R, which regulates calcium homeostasis and bone development^55^. PTH and PTHrP are equipotent agonists of the PTH1R, but the endocrine hormone PTH exhibits a slower PTH1R off-rate and longer duration cAMP signaling than the paracrine factor PTHrP^40,56^. The basis for their different off-rates was mapped to a single residue difference near the N-terminus of the peptides that binds deep within the PTH1R TMD^56^, so despite the conceptual similarities, PTH and AM2/IMD use very different structural mechanisms to attain their slow off-rates. The long receptor residence time of PTH allows it to continue to direct cAMP signaling from internalized PTH1R in endosomes, whereas PTHrP is limited to a traditional cAMP signal at the plasma membrane^40^. AM2/IMD might similarly direct AM_2_R signaling from endosomes because agonist stimulated AM_2_R undergoes internalization^57^, however, RAMP3 can interact with cytosolic factors that alter AM_2_R trafficking, in some cases by blocking internalization^58,59^. It will be important in the future to define how the long residence time of AM2/IMD affects AM_2_R signaling at the plasma membrane and inside the cell in diverse cell types, and to determine the biological functions of the AM2/IMD-AM_2_R cognate pair. These endeavors and future work to develop novel research tools and therapeutics targeting the AM receptors will benefit from the foundation laid here by revealing the mechanistic basis of the distinct AM2/IMD-AM_2_R binding kinetics.

## Materials and Methods

### Cell culture

COS-7 (African green monkey kidney fibroblast-like cell line, male; CRL 1651) and HEK293 (Human embryonic kidney cell line, female; CRL 1573) cells were from American Type Culture Collection (Manassas, VA, USA). Cells were cultured in Dulbecco’s modified Eagle’s medium (DMEM with 4.5 g/L glucose and L-glutamine) from Lonza (Basel, Switzerland) with 10 % v/v fetal bovine serum (Gibco 16000-044). Cells were grown at 37°C, 5% CO2 in a humidified incubator and passaged twice per week.

### Plasmid constructs

The human CLR and RAMPs were used throughout. N-terminally Nanoluciferase (NLuc)-tagged CLR was constructed in the pcDNA3.1(+) backbone using standard PCR and restriction enzyme cloning methods. The fusion construct was inserted between the EcoRI and KpnI sites and included a secretory signal peptide taken from the pHLsec vector^60^ followed by NLuc, a short linker, and residues 23-461 of CLR. The RAMP2/3 chimeras and the RAMP3 point mutants were ordered as synthetic GeneArt strings (Thermofisher) using the unoptimized human gene sequences including their natural signal sequences and with addition of EcoRI and XhoI restriction sites. The RAMP2/3 chimeras and RAMP3 mutants were inserted into the pcDNA3.1(+) vector using either restriction enzyme or Gibson assembly methods. All coding sequences were verified by DNA sequencing at the OUHSC laboratory of molecular biology and cytometry core facility. Protein sequences for the coding regions of each of the plasmids are provided in Table S5. The wild-type CLR and RAMP1-3 expression constructs in the pcDNA3.1(+) vector were from the cDNA resource center. The CAMYEL biosensor plasmid^36^ was a kind gift from Drs. Denise Wootten and Patrick Sexton.

### Synthetic peptides

Synthetic human peptides αCGRP(1-37), AM(13-52), and AM2/IMD(1-47) were purchased from Bachem (Bubendorf, Switzerland). Custom peptides were synthesized and HPLC-purified by Vivitide (Gardner, MA) or RS Synthesis (Louisville, KY). The lyophilized powders were resuspended at 5-10 mg/ml in sterile ultrapure water. DMSO was included at 9.5% v/v for AM(13-38)-AM2/IMD(34-47), 12.5% v/v for AM-TAMRA, and 10% v/v for AM2/IMD-TAMRA to improve solubility. Concentrations of the unlabeled peptides were determined by UV absorbance at 280 nm with dilutions in 10 mM Tris-HCl, 1 mM EDTA at pH 8.0. Extinction coefficients were calculated from Tyr, Trp, and cystine content. The concentration of CGRP was determined by the peptide content reported from Bachem. For the TAMRA-labeled peptides, concentrations were determined by visible absorbance at 560 nm with dilutions in 8M urea, 10 mM Tris-HCl, 1 mM EDTA, pH 8.0 and using the extinction coefficient of TAMRA (95,000 M^-1^cm^-1^). Peptides were stored as multiple aliquots at −80°C to limit the number of freeze-thaws. Table S6 lists the peptide sequences.

### Real-time BRET cAMP biosensor assay

COS-7 or HEK293 cells were seeded at 20,000 cells/well and 100 μL/well in a 96-well white plate and incubated for 24 hrs. The cells were transiently transfected with 250 ng total DNA and 375 ng of branched polyethylenimine (PEI) per well. For COS-7, 25 ng receptor, 25 ng RAMP, 125 ng CAMYEL biosensor, and 75 ng empty pcDNA3.1(+) was used. For HEK293, 10 ng receptor, 10 ng RAMP, 175 ng CAMYEL biosensor, and 55 ng empty vector was used. Two days after transfection the cells were washed with PBS and incubated for 30 min at room temperature in sterile filtered 25 mM NaHEPES pH 7.4, 104 mM NaCl, 5 mM KCl, 1 mM KH_2_PO_4_, 1.2 mM MgSO_4_, 2 mM CaCl_2_, 1 mg/mL fatty-acid-free bovine serum albumin (FAF-BSA), and 5 mM glucose. Coelenterazine h was added at 10 μM and incubated for 5 min at room temperature. The emissions at 475 and 535 nm were read in a PolarSTAR Omega plate reader (BMG Labtech, Ortenberg, Germany) using a dual luminescence optic for 5 min to establish the baseline. Agonist was manually added to 100 nM with reading for 15 min followed by manual addition of antagonist (10 μM) or forskolin (10 μM) or buffer with an additional 40 min reading. The antagonists used were αCGRP(8-37) [N31D/S34P/K35W/A36S] for CLR-RAMP1 and AM(22-52) [S48G/Q50W] for CLR-RAMP2/3. For assays performed at 37°C, the cells were incubated in a 37°C incubator and the plate reader was heated to 37°C. The 475/535 BRET ratio was used to plot the data over time with buffer control subtracted from each curve. The decay phase after antagonist addition was fit to a one-phase exponential decay model in GraphPad Prism v9.4.1 (GraphPad Software).

### LANCE cAMP accumulation assay

cAMP accumulation assays were performed as previously described^31–33^. In brief, COS-7 cells were seeded in a 96-well plate on day one. Cells were transiently transfected with receptors on day two and on day four the cells were stimulated with agonist for 15 min at 37°C in the presence of IBMX. The cells were lysed and the cAMP was measured using a LANCE *ultra* cAMP Detection Kit (Perkin Elmer) according to the manufacturer’s instructions.

### NLuc-CLR-RAMP3 membranes preparation

COS-7 cells were seeded at 2.5 million cells/dish into ten 150 mm^2^ plastic culture dishes and grown for 2 days. The cells were transiently transfected with 50 μg DNA (2 μg NLuc-CLR, 2 μg RAMP3, and 46 μg empty pcDNA3.1) and 75 μg PEI/dish. After two days the media was aspirated and the cells were washed with PBS and harvested with ice-cold PBS and 5 mM EDTA with a cell scraper and pelleted in a centrifuge at 1000 xg for 5 min at 4°C. All remaining steps were on ice or at 4°C. The pellets were resuspended in hypotonic buffer (25 mM NaHEPES pH 7.5, 2 mM MgCl_2_, 1 mM EDTA with 1X EDTA-free PIERCE protease inhibitor tablet (PI)). The cells were homogenized with an Ultra Turrax for 30 s at 10k rpm followed by a 10 min incubation. Cell debris was pelleted at 800 xg for 10 min. The supernatant was transferred to ultracentrifuge tubes and centrifuged at 100k xg for 1 hr. The cell pellets were resuspended in wash buffer (25 mM NaHEPES pH 7.5, 250 mM NaCl, 2 mM MgCl_2_, 1 mM EDTA, and 1X PI). The membranes were homogenized and spun in the ultracentrifuge as before. The membrane pellets were combined in storage buffer (25 mM NaHEPES pH 7.5, 25 mM NaCl, 2 mM MgCl_2_, 10 % v/v glycerol, and 1X PI), homogenized, aliquoted, and flash-frozen in liquid nitrogen for storage at −80°C The protein concentration was determined using the DC protein assay (BioRad) per the manufacturer’s instructions.

### nanoBRET ligand binding assays

Each binding assay format used NLuc-CLR-RAMP3 membranes at a concentration of 0.0075 mg/mL in a binding buffer of 25 mM NaHEPES pH 7.4, 104 mM NaCl, 5 mM KCl, 1 mM KH_2_PO_4_, 3 mM MgSO_4_, 2 mM CaCl_2_, 1 mg/mL FAF-BSA, 50 μg/mL saponin, and 50 μM GTPγS (uncoupled state) or 30 μM H_6_-sumo-miniGs^17^ (coupled state). The equilibrium assays were performed at room temperature and the kinetic assays were at 25°C. For equilibrium binding, 3-fold serial dilutions of AM-TAMRA and AM2/IMD-TAMRA were incubated with the membranes for 3hr (uncoupled state) or 4.5hr (coupled state) at room temperature in a 96-well white plate. Furimazine substrate (Promega) was added at 1X and incubated for 5 min. Emission was read with 460 nm and 610 long pass filters in the PolarSTAR Omega plate reader. Agonist concentration was plotted against the BRET ratio of 610/460 and fit to a one site specific binding model in GraphPad Prism after subtracting the buffer control.

For association kinetics, 2-fold serial dilutions of AM-TAMRA and AM2/IMD-TAMRA in binding buffer were added to a 96 well white plate at 2X. The membranes at 2X were incubated for 15 min in binding buffer with GTPγS followed by addition of furimazine at 2X with a further 5 min incubation and then loaded into the plate reader injector. The membranes were injected into the peptide serial dilutions in the plate with reading every 16 s for 40 min. The BRET ratio 610/460 was plotted against time with background subtraction and each curve was fit to a one-phase association exponential model in GraphPad Prism using the first 10 min of data to minimize the effects of signal decay.

For dissociation kinetics, the membranes and AM-TAMRA or AM2/IMD-TAMRA (10 nM) in binding buffer were incubated in a 96-well white plate for 15 min followed by furimazine addition at 1X for 5 min. Emission at 460 and 610 nm were read every 13 sec for 5 min to establish a baseline prior to injection of buffer or antagonist to initiate dissociation. Binding buffer or AM(22-52) [S48G/Q50W] antagonist (1 μM) loaded into the injectors were injected into the plate, which was read for 1-2 hr. The BRET ratio 610/460 was plotted against time with background subtraction. The curves were normalized to their corresponding buffer control injections as 100% to account for the signal decay with time. The normalized dissociation curves were fit to a two-phase exponential decay model in GraphPad Prism constraining the AM-TAMRA and AM2/IMD-TAMRA curves to have the same plateau.

### Native PAGE assays for agonist-GPCR-miniGs coupling and thermostability

The coupling assays were performed as previously described up to the point of formation of quaternary agonist-CLR-RAMP3-miniGs complexes, and they used the previous MBP-CLR-EGFP:MBP-RAMP3 membrane preparation and H_6_-sumo-miniGs freshly purified as described^17,43^. Three-fold serial dilutions of AM(22-52) [S48G/Q50W] antagonist were added to the pre-formed quaternary complexes and incubated for an additional 2 or 19 hours at 4°C on a rocking shaker. The samples were centrifuged, and the supernatants were analyzed by native PAGE with EGFP fluorescence imaging as described^17,43^. The thermostability assays were performed as previously described using LMNG/CHS^17^, except with the inclusion of ligands and using finer temperature increments.

### Rebuilding and refinement of cryo-EM structures in preparation for MD

AlphaFold2^46^ was used via Colabfold^61^ to predict the structures of the AM- and AM2/IMD-bound CLR-RAMP3 complexes. Using Pymol (Schrodinger, LLC) and/or Coot^62^, the receptor ECD complexes from the AlphaFold models were extracted and the peptides were modeled with the AM and AM2/IMD conformations from high-resolution crystal structures of their complexes with CLR-RAMP2 ECD (4RWF) and CLR-RAMP1 ECD (6D1U), respectively. These peptide-bound CLR-RAMP3 ECD complex models were used to replace the corresponding complexes in 6UUS (AM) and 6UVA (AM2/IMD) as starting points for rebuilding. The models were manually rebuilt in Coot guided by fitting to the multiple cryo-EM density maps deposited in the EMDB for the AM (20901) and AM2/IMD (20906) structures. The “combine focused maps” tool in Phenix^63^ was used to generate a single best composite map from the multiple maps for each structure and the rebuilt models were refined to their respective composite maps using Phenix real-space refinement.

### MD simulations

The rebuilt structures were prepared for MD simulation by rebuilding two missing CLR loops with MODELLER^64^: 351-362 and 323-329. The membrane-solvated system was then constructed in CHARMM-GUI^65^, retaining the two resolved cholesterol molecules. In the CHARMM-GUI system preparation tool, GlcNAc glycosylations were added to N66, N118 and N123 of CLR, as well as to N29, N58, N71 and N103 of RAMP3. The C-terminal residue of each peptide was amidated using the CT2 patch. The membranes were constructed using a roughly 3:2 ratio of POPC:cholesterol, with a 10 Å clearance of the protein towards the edge of the box in the x and y dimensions. This resulted in 27 cholesterol and 39 POPC molecules in the lower (intracellular) leaflet and 26 cholesterol and 39 POPC molecules in the upper (extracellular) leaflet for the AM-AM_2_R system. The AM2/IMD-AM_2_R system was slightly smaller in the x and y dimensions, resulting in 26 cholesterol and 36 POPC in the lower leaflet and 24 cholesterol and 36 POPC in the upper leaflet.

The systems were solvated using a 10 Å clearance with TIP3 water molecules, resulting in 16029 and 15559 waters for the AM-AM_2_R and AM2/IMD-AM_2_R systems, respectively. NaCl was added to each system at a concentration of 0.15 M. The equilibration and heating steps were run as specified in the scripts provided by CHARMM-GUI, with the addition of an auxiliary restraint force acting on all non-hydrogen CLR atoms with a z-coordinate greater than 9.5 nm. This force was meant to mimic the stabilizing effect of the G protein, even though it was omitted from the simulation to minimize the computational cost. The restraint force was implemented using a CustomExternalForce function in OpenMM^66^, that restrained the positions of the atoms to their original positions using a harmonic restraint energy with a force constant of 5000 kJ/mol/nm^2^. This force acted on the entire C-terminal helix of CLR (residues 389 to 402), as well as residues from the other three intracellular loops (residues 165-172, 240-251, 318-328). The equilibration included 5000 steps of energy minimization followed by a six-step minimization protocol (125000 or 250000 steps each, 1 fs or 2 fs timestep), where positional restraints on the protein backbone, protein side-chain and lipids, were gradually reduced. Dihedral restraints on lipids and carbohydrates were similarly reduced over the course of the equilibration. An OpenMM MonteCarloMembraneBarostat function was employed, with a pressure coupling frequency of 100 steps and a reference pressure of 1 bar. Following equilibration, all restraints besides the G protein CLR restraints were removed and the system was allowed to relax for 10 ns at a 2 fs timestep.

The final frames of each system were used as initial structures for a set of weighted ensemble simulations with the REVO (Resampling Ensembles by Variation Optimization) method^48^ using the wepy software^67^. Three independent replicates, with 48 trajectories each, were used for each system. Each ensemble was run for 2000 cycles of resampling, with 20 ps of dynamics in each cycle. The combined sampling time across all replicates was 5.76 *μ*s per system, or 11.52 *μ*s in total. The resampling procedure following each cycle used the REVO resampling function, with a minimum probability of 10^-12^, a maximum probability of 0.1 and a weight-based novelty function with a distance exponent of 4, following previous work^48,68^. The distance metric used for the trajectory variation function was the RMSD of the peptide following alignment to the entire CLR protein. This distance is calculated between the trajectories in the ensemble and is used to identify “outliers” that are preferentially chosen for cloning, as well as quasi-redundant trajectories that are preferentially chosen for merging. For more information on the REVO algorithm, please refer to the following references^48^. Merging operations could only be performed for trajectories that were within 8 Å of each other, according to the distance metric described above. The vast majority of the trajectory pairs met this criterion.

All probability-based analyses of these datasets, such as the RMSD probability distributions and distance probability distributions, employed the weights of the trajectories. Water density isosurfaces were computed using the volmap tool of VMD^69^.

### Statistical analysis

For all assays, duplicate technical replicates were used, except for the native PAGE assays, and reported as mean ± SD in the representative plots. All experiments were done with three independent replicates on different days and reported as mean ± SEM. Statistical analysis was performed using GraphPad Prism on the log form of the values. One-way ANOVA with the Tukey’s post hoc test was used at a confidence interval of 99.9% reaching a statistical significance of p<0.001, comparing the mean of the three independent replicates. Only the most important comparisons that reached statistical significance were shown in figures.

## Acknowledgments

We thank Drs. Denise Wootten and Patrick Sexton for kindly providing the CAMYEL plasmid, and Dr. Amanda Roehrkasse for providing the receptor cartoons used in Figure 3A and Figure 8.

## Funding

National Institutes of Health grant R01GM104251 (AAP)

National Institutes of Health grant R01GM130794 (ARD)

National Science Foundation grant DMS1761320 (ARD)

## Author contributions

Conceptualization: KMB, AAP

Methodology: KMB, ARD, AAP

Investigation: KMB, JAK, PHG, JL, ARD, AAP

Supervision: ARD, AAP

Visualization: KMB, ARD, AAP

Writing—original draft: KMB, ARD, AAP

Writing—review & editing: KMB, ARD, AAP

## Competing interests

KMB and AAP are filing a patent on kinetically selective AM and AM2/IMD variants.

