## Supplemental Figures and Tables for "Adrenomedullin 2/intermedin is a slow off-rate, long-acting endogenous agonist of the adrenomedullin_2_ G protein-coupled receptor"

**This file includes:**

Figs. S1 to S5

Tables S1 to S6

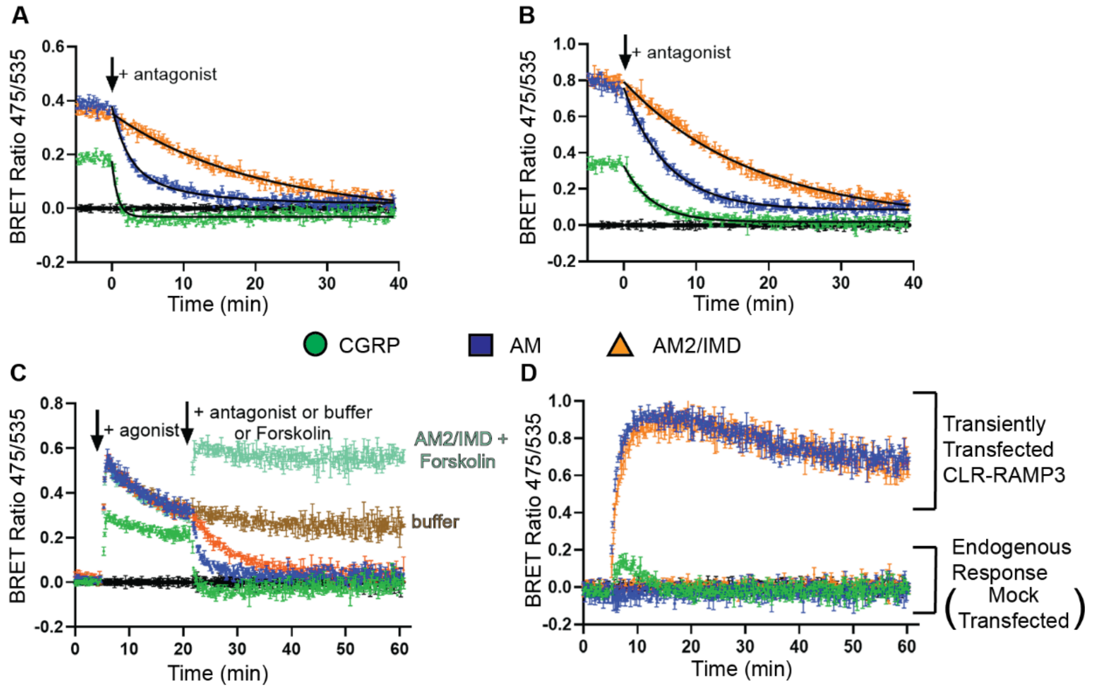


**Fig. S1. CGRP, AM, and AM2/IMD cAMP signaling kinetics controls.**

**(A, B)** One-phase exponential decay curve fit for CLR-RAMP3 after antagonist addition in COS-7 (A) and HEK293 (B) cells. **(C)** cAMP kinetics for CLR-RAMP3 in COS-7 cells at 37°C with CGRP (green circle), AM (blue square), or AM2/IMD (orange triangle) used as agonists and 10 µM AM(22-52) [S48G/Q50W] as the antagonist. The brown and cyan curves indicated AM2/IMD stimulation followed by buffer addition or 10 µM forskolin control, respectively. **(D)** Endogenous response to 100 nM CGRP, AM, and AM2/IMD in HEK293 cells compared to transiently transfected CLR-RAMP3 response to 100 nM AM and AM2/IMD.

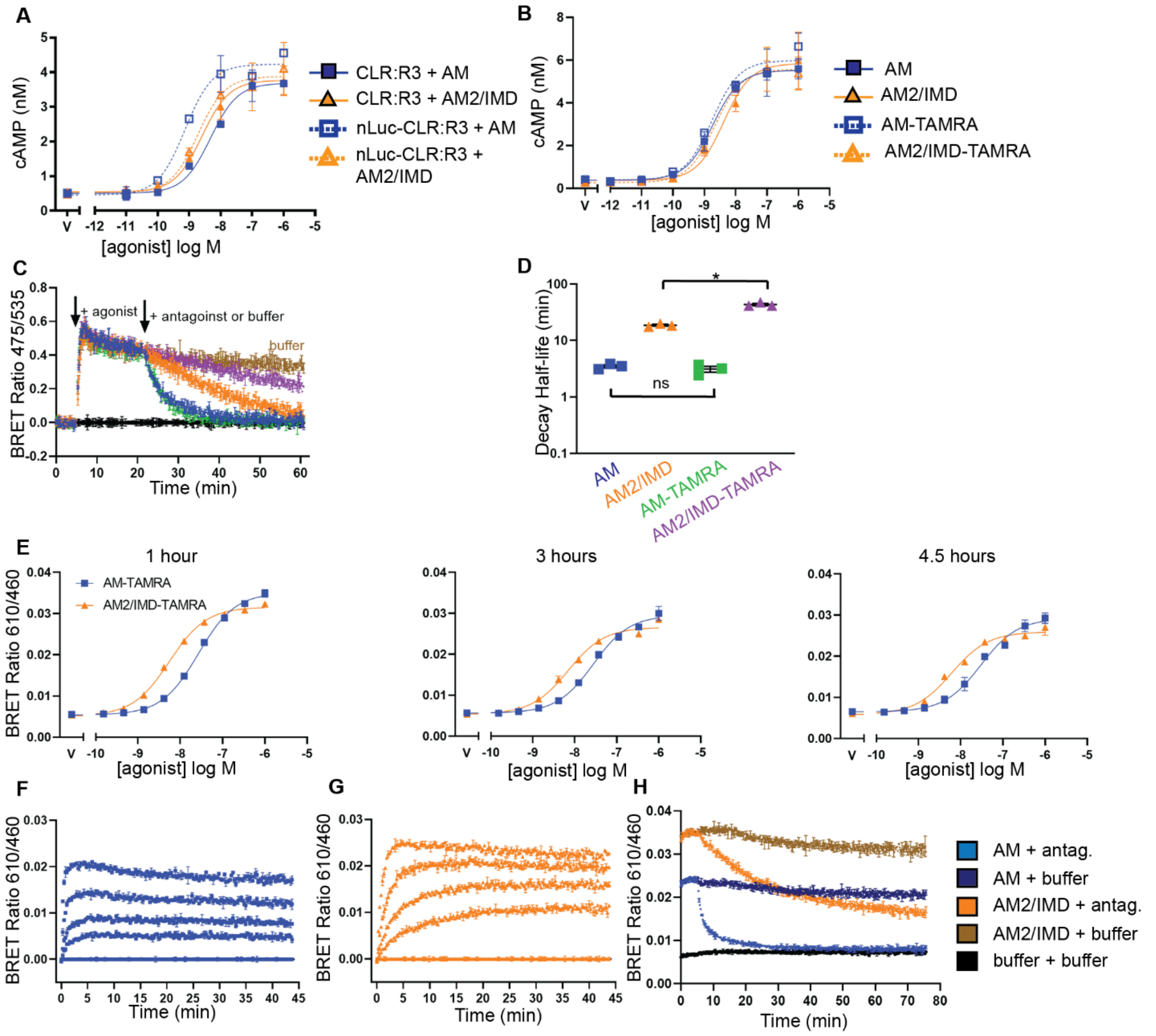


**Fig. S2. Controls for nanoBRET binding kinetics of AM-TAMRA and AM2/IMD-TAMRA to NLuc-CLR:RAMP3 membranes.**

**(A)** cAMP accumulation assay for the indicated receptor-agonist combinations in COS-7 cells. **(B)** cAMP accumulation assay for the indicated peptides at CLR-RAMP3 in COS-7 cells. **(C)** cAMP signaling kinetics at CLR-RAMP3 in COS-7 cells with 100 nM of the indicated agonist and 10 µM AM(22-52) [S48G/Q50W] antagonist. **(D)** Scatter plot summarizing the decay half-lives from panel C with mean ± SEM of three independent replicates. AM-TAMRA had a half-life of 3.1 ± 0.65 min and AM2/IMD-TAMRA had a half-life of 43 ± 3.5 min. Statistical analysis done with one-way ANOVA and Tukey's post hoc test. **(E)** Equilibrium binding with the indicated incubations times. **(F, G)** Association kinetics of AM-TAMRA corresponding to Fig. 2D with extended time (F) and AM2/IMD-TAMRA corresponding to Fig. 2E (G). **(H)** Raw dissociation kinetic data corresponding to Fig. 2F with AM-TAMRA (blue) and AM2/IMD-TAMRA (orange) showing slight signaling decay over time.

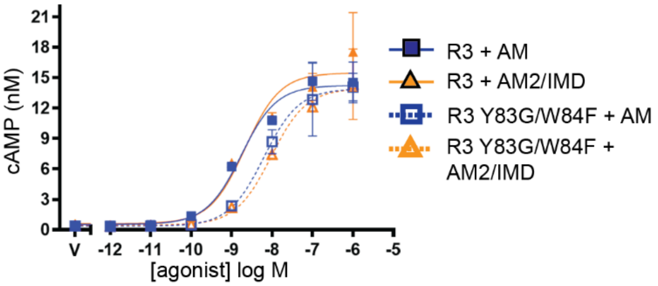


**Fig. S3. cAMP accumulation assay for AM and AM2/IMD with CLR-RAMP3 wild-type and CLR-RAMP3 Y83G/W84F.**

CLR was co-expressed with WT RAMP3 or RAMP3 Y83G/W84F and stimulated with AM or AM2/IMD in COS-7 cells. Values plotted as mean ± SD of technical replicates show as a representative of two independent replicates.

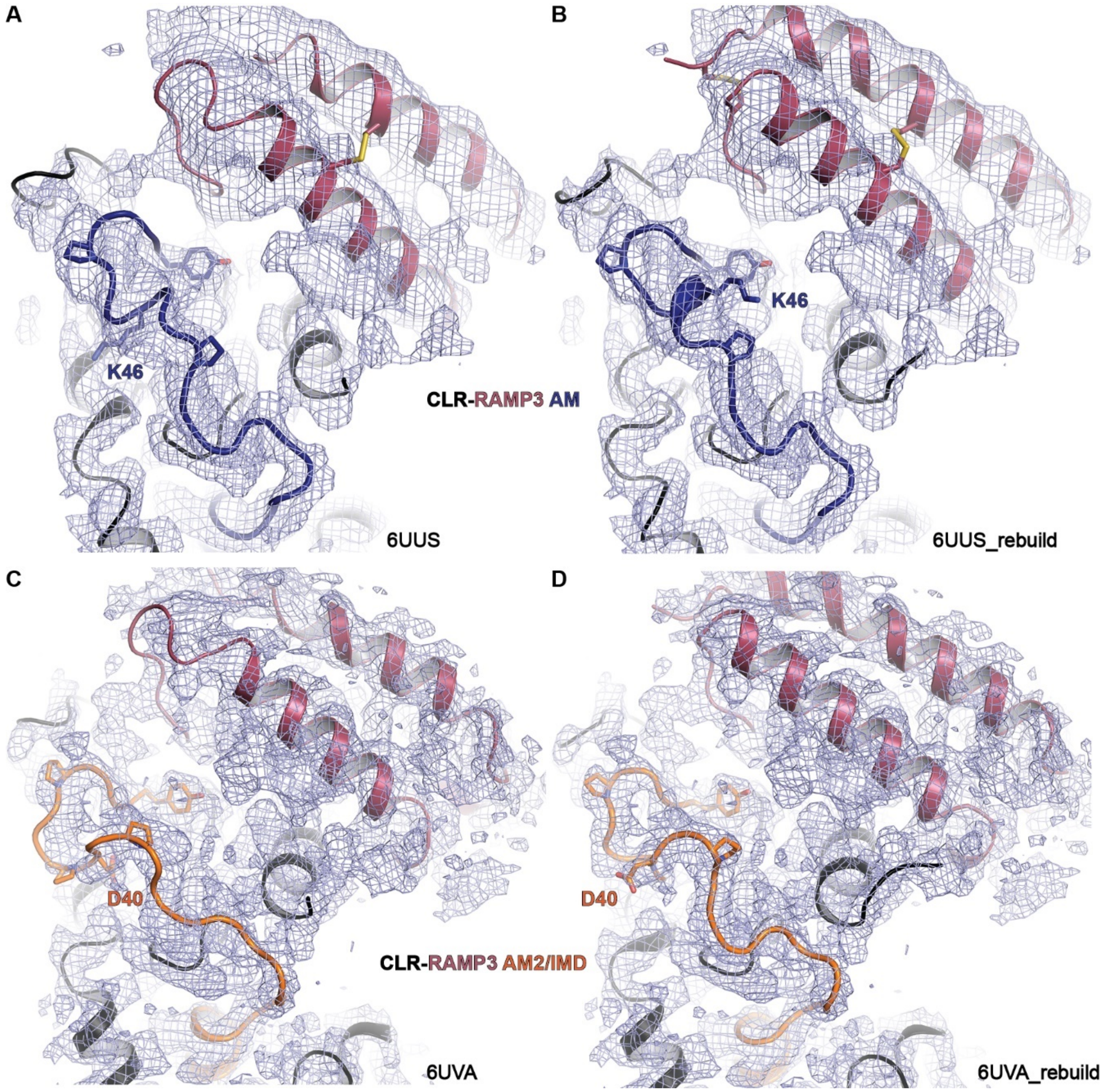


**Fig. S4. Rebuilding of the AM- and AM2/IMD-bound CLR-RAMP3 cryo-EM structures.** (A) AM-AM$_2$R-Gs (6UUS). (B) Rebuilt and refined 6UUS. (C) AM2/IMD-AM$_2$R-Gs (6UVA). (D) Rebuilt and refined 6UVA. The structures are shown along with their respective composite cryo-EM density maps generated as described in Materials and Methods.

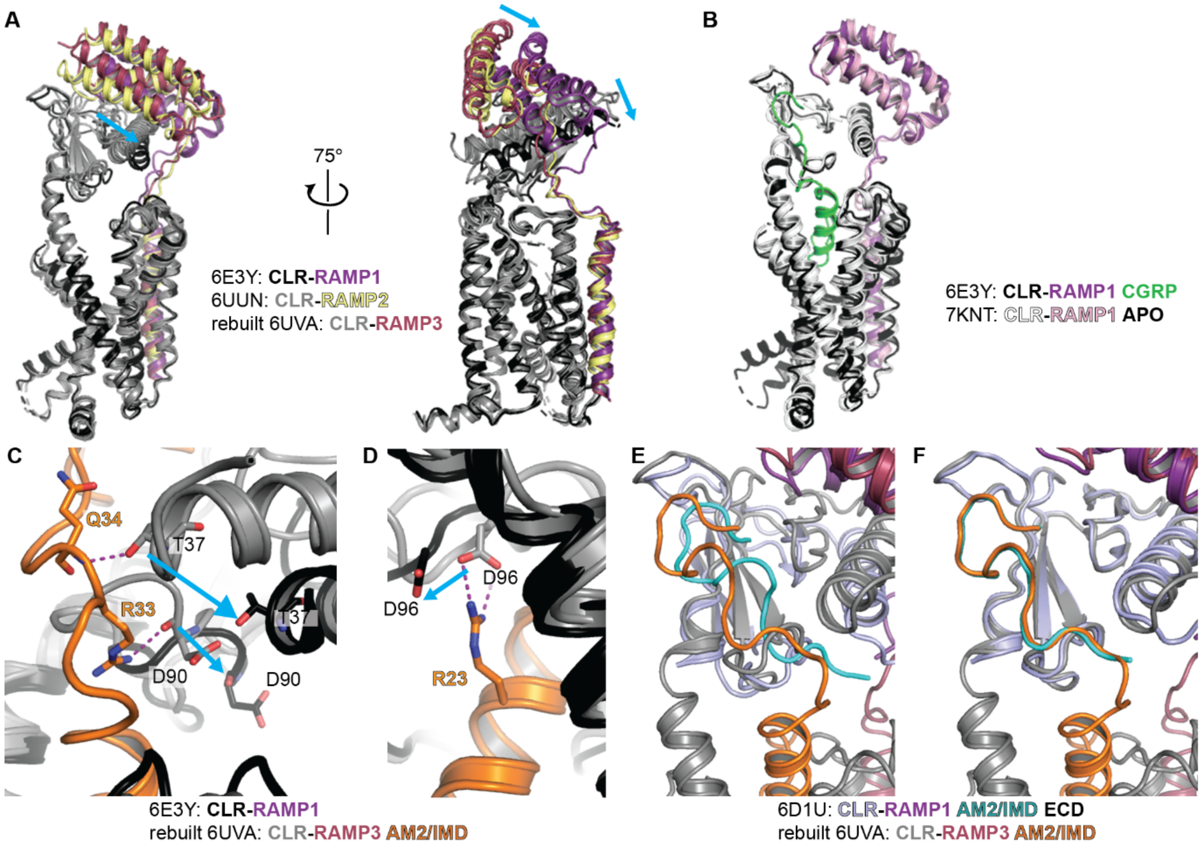


**Fig. S5. Structural comparisons of CLR ECD-TMD arrangements in CLR-RAMP heterodimers.**

**(A)** Overlay of the cryo-EM structures of the indicated CLR-RAMP complexes aligned to the CLR TMD. Arrows depict the shift in the ECD positions of the RAMP1 complex as compared to the RAMP2/3 complexes. For clarity, the peptides are not shown. **(B)** Overlay of the indicated APO and CGRP-bound CLR-RAMP1 cryo-EM structures showing that RAMP1 is responsible for the CLR ECD-TMD arrangement rather than the peptide. **(C, D)** The panels depict an overlay of the indicated structures aligned to the CLR TMD. For clarity, the CGRP peptide is not shown. Depiction of the CLR residues T37, D90, and D96 that contact the AM2/IMD mid-region, showing these key interactions between AM2/IMD and CLR-RAMP3 are incompatible with the CLR ECD-TMD arrangement induced by RAMP1. **(E)** Rebuilt 6UVA overlaid with the AM2/IMD-bound CLR-RAMP1 ECD crystal structure (6D1U) in the same position observed in the cryo-EM structure of full-length CLR-RAMP1 (6E3Y). This shows that the altered CLR ECD-TMD arrangement induced by RAMP1 is incompatible with full-length AM2/IMD adopting the same conformation observed in rebuilt 6UVA (RAMP3). **(F)** As in E except that the AM2/IMD-bound CLR-RAMP1 ECD crystal structure was aligned to the CLR ECD in rebuilt 6UVA. This shows that the ECD-binding segment of AM2/IMD adopts the same conformation in the RAMP1 and RAMP3 complexes.

**Table S1. Summary of cAMP signaling decay kinetics for CGRP, AM, and AM2/IMD in COS-7 and HEK293 cells at each receptor heterodimer**

| | COS-7 | | | HEK293 | | |
|---|---|---|---|---|---|---|
| | Observed decay rate ($min^{-1}$ ± SEM) | Time Constant τ (min ± SEM)* | Half-life (min ± SEM)† | Observed decay rate ($min^{-1}$ ± SEM) | Time Constant τ (min ± SEM) | Half-life (min ± SEM) |
| CLR-RAMP1 | | | | | | |
| CGRP | 0.26 ± 0.0024 | 3.80 ± 0.035 | 2.6 ± 0.024 | 0.16 ± 0.028 | 6.7 ± 1.0 | 4.6 ± 0.70 |
| AM | 0.87 ± 0.0055 | 1.2 ± 0.0073 | 0.80 ± 0.0051 | 0.22 ± 0.035 | 4.8 ± 0.67 | 3.3 ± 0.46 |
| AM2/IMD | 0.49 ± 0.038 | 2.1 ± 0.15 | 1.4 ± 0.10 | 0.21 ± 0.027 | 4.9 ± 0.57 | 3.4 ± 0.39 |
| CLR-RAMP2 | | | | | | |
| CGRP | 0.80 ± 0.042 | 1.3 ± 0.065 | 0.87 ± 0.045 | 0.27 ± 0.0077 | 3.8 ± 0.11 | 2.6 ± 0.073 |
| AM | 0.33 ± 0.025 | 3.1 ± 0.22 | 2.1 ± 0.15 | 0.19 ± 0.040 | 5.8 ± 1.1 | 4.0 ± 0.76 |
| AM2/IMD | 0.50 ± 0.025 | 2.0 ± 0.11 | 1.4 ± 0.074 | 0.22 ± 0.018 | 4.6 ± 0.37 | 3.2 ± 0.25 |
| CLR-RAMP3 | | | | | | |
| CGRP | 1.2 ± 0.029 | 0.85 ± 0.022 | 0.59 ± 0.015 | 0.31 ± 0.053 | 3.5 ± 0.51 | 2.4 ± 0.35 |
| AM | 0.21 ± 0.031 | 4.9 ± 0.67 | 3.4 ± 0.46 | 0.19 ± 0.042 | 5.9 ± 1.2 | 4.1 ± 0.80 |
| AM2/IMD | 0.045 ± 0.0075 | 24 ± 4.7 | 17 ± 3.3 | 0.066 ± 0.022 | 19 ± 5.5 | 13 ± 3.8 |

* Time constant calculated as the inverse of decay rate
† Half-life calculated as ln 2 divided by the decay rate

**Table S2. Summary of nanoBRET binding parameters for Nluc-CLR:RAMP3 membranes.**

| | | + GTPγS | | + miniGs | |
|---|---|---|---|---|---|
| | | AM-TAMRA | AM2/IMD-TAMRA | AM-TAMRA | AM2/IMD-TAMRA |
| Equilibrium Binding | Kd (nM) | 26 ± 1.6 | 6.8 ± 0.81 | 8.0 ± 0.68 | 2.9 ± 0.18 |
| Association Kinetics | $k_{on}$ ($M^{-1}min^{-1}$ ± SEM)* | $1.4x10^8$ ± $2.3x10^6$ | $4.1x10^7$ ± $2.6x10^6$ | ND† | ND |
| | $k_{off}$ ($min^{-1}$ ± SEM)‡ | 0.28 ± 0.027 | 0.032 ± 0.030 | ND | ND |
| | Calculated Kd (nM ± SEM)§ | 2.0 ± 0.096 | 0.79 ± 0.093 | ND | ND |
| | Time Constant τ (min ± SEM)** | 3.6 ± 0.095 | 31 ± 0.93 | ND | ND |
| | Half-life (min ± SEM)†† | 2.5 ± 0.066 | 21 ± 0.64 | ND | ND |
| Dissociation Kinetics | $k_{off}$ fast ($min^{-1}$ ± SEM) | 0.93 ± 0.020 | 0.099 ± 0.0020 | 0.15 ± 0.0044 | 0.044 ± 0.0008 |
| | Time Constant τ fast (min ± SEM) | 1.1 ± 0.023 | 10 ± 0.20 | 6.8 ± 0.21 | 23 ± 0.40 |
| | Half-life fast (min ± SEM) | 0.75 ± 0.016 | 7.0 ± 0.14 | 4.7 ± 0.14 | 16 ± 0.28 |
| | $k_{off}$ slow ($min^{-1}$ ± SEM) | 0.10 ± 0.0031 | 0.00091 ± 0.00020 | 0.0064 ± 0.00005 | 0.0022 ± 0.00006 |
| | Time Constant τ slow (min ± SEM) | 9.9 ± 0.31 | 110 ± 2.4 | 157 ± 1.2 | 456 ± 13 |
| | Half-life slow (min ± SEM) | 6.9 ± 0.21 | 76 ± 1.7 | 109 ± 0.81 | 316 ± 8.9 |
| | Percent Fast (% ± SEM) | 66.5 ± 1.2 | 34.7 ± 0.87 | 61 ± 0.93 | 22 ± 0.55 |

* Values obtained from the slope in linear regression shown in Fig. 2E
† ND = not done
‡ Values obtained from the y-intercept in linear regression shown in Fig. 2E
§ Calculated as off-rate/on-rate
** Calculated as the inverse of the off-rate
†† Calculated by ln 2 divided by the off-rate

### Table S3. Summary of chimeric peptide cAMP signaling decay kinetic values

| | | Observed decay rate ($min^{-1}$ ± SEM) | Time Constant τ (min ± SEM)[*] | Half-life (min ± SEM)[†] |
|---|---|---|---|---|
| CLR-RAMP1 | AM | 0.92 ± 0.033 | 1.1 ± 0.039 | 0.76 ± 0.027 |
| | AM2/IMD | 0.50 ± 0.018 | 2.0 ± 0.075 | 1.4 ± 0.052 |
| | AM-AM2 half | 0.73 ± 0.021 | 1.4 ± 0.038 | 0.96 ± 0.027 |
| | AM2-AM half | 0.82 ± 0.027 | 1.2 ± 0.039 | 0.85 ± 0.027 |
| | AM-AM2 ECD | 0.72 ± 0.041 | 1.4 ± 0.086 | 0.98 ± 0.060 |
| | AM2-AM ECD | 0.48 ± 0.031 | 2.1 ± 0.13 | 1.4 ± 0.089 |
| | AM-AM2-AM | 0.70 ± 0.034 | 1.4 ± 0.074 | 1.0 ± 0.051 |
| | AM2-AM-AM2 | 0.89 ± 0.081 | 1.2 ± 0.099 | 0.80 ± 0.069 |
| CLR-RAMP2 | AM | 0.33 ± 0.020 | 3.1 ± 0.19 | 2.2 ± 0.13 |
| | AM2/IMD | 0.51 ± 0.011 | 2.0 ± 0.042 | 1.4 ± 0.029 |
| | AM-AM2 half | 0.81 ± 0.021 | 1.2 ± 0.031 | 0.86 ± 0.021 |
| | AM2-AM half | 0.53 ± 0.035 | 1.9 ± 0.13 | 1.3 ± 0.092 |
| | AM-AM2 ECD | 0.60 ± 0.025 | 1.7 ± 0.074 | 1.2 ± 0.051 |
| | AM2-AM ECD | 0.12 ± 0.0056 | 8.7 ± 0.41 | 6.0 ± 0.29 |
| | AM-AM2-AM | 0.042 ± 0.0048 | 25 ± 3.0 | 17 ± 2.1 |
| | AM2-AM-AM2 | 0.86 ± 0.065 | 1.2 ± 0.089 | 0.82 ± 0.061 |
| CLR-RAMP3 | AM | 0.22 ± 0.019 | 4.8 ± 0.42 | 3.3 ± 0.29 |
| | AM2/IMD | 0.039 ± 0.0030 | 27 ± 1.8 | 19 ± 1.3 |
| | AM-AM2 half | 0.13 ± 0.011 | 8.0 ± 0.72 | 5.6 ± 0.50 |
| | AM2-AM half | 0.19 ± 0.016 | 5.5 ± 0.50 | 3.8 ± 0.35 |
| | AM-AM2 ECD | 0.14 ± 0.011 | 7.0 ± 0.56 | 4.9 ± 0.39 |
| | AM2-AM ECD | 0.022 ± 0.0031 | 48 ± 7.3 | 33 ± 5.0 |
| | AM-AM2-AM | 0.015 ± 0.0015 | 66 ± 6.0 | 49 ± 4.2 |
| | AM2-AM-AM2 | 0.60 ± 0.049 | 1.7 ± 0.14 | 1.2 ± 0.099 |

[*] Time constant calculated as inverse of decay rate
[†] Half-life calculated as ln 2 divided by decay rate

### Table S4. Summary of cAMP signaling decay kinetic values for RAMP2/3 chimeras and RAMP3 mutants

| | AM | | | AM2/IMD | | |
|---|---|---|---|---|---|---|
| | Observed decay rate ($min^{-1}$ ± SEM) | Time Constant τ (min ± SEM)* | Half-life (min ± SEM)† | Observed decay rate ($min^{-1}$ ± SEM) | Time Constant τ (min ± SEM) | Half-life (min ± SEM) |
| RAMP Chimeras | | | | | | |
| RAMP2 | 0.34 ± 0.020 | 3.1 ± 0.13 | 2.1 ± 0.11 | 0.49 ± 0.036 | 2.1 ± 0.079 | 1.4 ± 0.097 |
| RAMP3 | 0.18 ± 0.011 | 5.5 ± 0.17 | 3.9 ± 0.25 | 0.035 ± 0.0047 | 29 ± 3.0 | 21 ± 3.3 |
| R2wR3 ECD | 0.22 ± 0.28 | 4.7 ± 0.53 | 3.2 ± 0.37 | 0.053 ± 0.0095 | 33 ± 1.7 | 23 ± 1.2 |
| R3wR2 ECD | 0.23 ± 0.016 | 4.3 ± 0.28 | 3.0 ± 0.19 | 0.36 ± 0.0038 | 2.8 ± 0.030 | 2.0 ± 0.021 |
| R2wR3 TMD | 0.27 ± 0.011 | 3.7 ± 0.16 | 2.6 ± 0.11 | 0.41 ± 0.015 | 2.4 ± 0.083 | 1.7 ± 0.058 |
| R3wR2 TMD | 0.16 ± 0.0066 | 6.3 ± 0.27 | 4.3 ± 0.19 | 0.037 ± 0.0084 | 30 ± 6.9 | 21 ± 4.8 |
| R2wR3 C-tail | 0.28 ± 0.026 | 3.7 ± 0.32 | 2.6 ± 0.22 | 0.54 ± 0.013 | 1.9 ± 0.044 | 1.3 ± 0.030 |
| R3wR2 C-tail | 0.22 ± 0.014 | 4.6 ± 0.30 | 3.2 ± 0.21 | 0.034 ± 0.0020 | 30 ± 1.8 | 21 ± 1.3 |
| RAMP3 Mutant | | | | | | |
| RAMP3 | 0.25 ± 0.013 | 4.0 ± 0.23 | 2.8 ± 0.16 | 0.048 ± 0.0038 | 22 ± 1.9 | 15 ± 1.3 |
| RAMP3 Y83G/W84F | 0.40 ± 0.012 | 2.5 ± 0.070 | 1.7 ± 0.049 | 0.23 ± 0.0081 | 4.3 ± 0.16 | 3.0 ± 0.11 |

* Time constant calculated as inverse decay rate
† Half-life calculated as ln 2 divided by decay rate

**Table S5. Amino acid sequences of plasmid constructs**

| Plasmid Number | Protein sequence |
|---|---|
| pKB003 (NLuc-CLR) | MGILPSPGMPALLSLVSLLSVLLMGCVAETGMVFTLEDFVGDWRQTAGYN LDQVLEQGGVSSLFQNLGVSVTPIQRIVLSGENGLKIDIHVIIPYEGLSGDQ MGQIEKIFKVVYPVDDHHFKVILHYGTLVIDGVTPNMIDYFGRPYEGIAVFD GKKITVTGTLWNGNKIIDERLINPDGSLLFRVTINGVTGWRLCERILAGSGIS ELEESPEDSIQLGVTRNKIMTAQYECYQKIMQDPIQQAEGVYCNRTWDGW LCWNDVAAGTESMQLCPDYFQDFDPSEKVTKICDQDGNWFRHPASNRT WTNYTQCNVNTHEKVKTALNLFYLTIIGHGLSIASLLISLGIFFYFKSLSCQRI TLHKNLFFSFVCNSVVTIIHLTAVANNQALVATNPVSCKVSQFIHLYLMGCN YFWMLCEGIYLHTLIVVAVFAEKQHLMWYYFLGWGFPLIPACIHAIARSLYY NDNCWISSDTHLLYIIHGPICAALLVNLFFLLNIVRVLITKLKVTHQAESNLYM KAVRATLILVPLLGIEFVLIPWRPEGKIAEEVYDYIMHILMHFQGLLVSTIFCF FNGEVQAILRRNWNQYKIQFGNSFSNSEALRSASYTVSTISDGPGYSHDC PSEHLNGKSIHDIENVLLKPENLYN |
| pKB016 (R2wR3 ECD) | METGALRRPQLLPLLLLLCGGCPRAGGCNETGMLERLPLCGKAFADMMG KVDVWKWCNLSEFIVYYESFTNCTEMEANVVGCYWPNPLAQGFITGIHRQ FFSNCSLVQPTFSDPPEDVLLAMIIAPICLIPFLITLVVWRSKDSEAQA |
| pKB015 (R3wR2 ECD) | MASLRVERAGGPRLPRTRVGRPAALRLLLLLGAVLNPHEALAQPLPTTGTP GSEGGTVKNYETAVQFCWNHYKDQMDPIEKDWCDWAMISRPYSTLRDCL EHFAELFDLGFPNPLAERIIFETHQIHFANCTVDRVHLEDPPDEVLIPLIVIPV VLTVAMAGLVVWRSKRTDTLL |
| pKB017 (R2wR3 TMD) | MASLRVERAGGPRLPRTRVGRPAALRLLLLLGAVLNPHEALAQPLPTTGTP GSEGGTVKNYETAVQFCWNHYKDQMDPIEKDWCDWAMISRPYSTLRDCL EHFAELFDLGFPNPLAERIIFETHQIHFANCSLVQPTFSDPPDEVLIPLIVIPV VLTVAMAGLVVWRSKDSEAQA |
| pKB018 (R3wR2 TMD) | METGALRRPQLLPLLLLLCGGCPRAGGCNETGMLERLPLCGKAFADMMG KVDVWKWCNLSEFIVYYESFTNCTEMEANVVGCYWPNPLAQGFITGIHRQ FFSNCTVDRVHLEDPPEDVLLAMIIAPICLIPFLITLVVWRSKRTDTLL |
| pKB019 (R2wR3 C-tail) | MASLRVERAGGPRLPRTRVGRPAALRLLLLLGAVLNPHEALAQPLPTTGTP GSEGGTVKNYETAVQFCWNHYKDQMDPIEKDWCDWAMISRPYSTLRDCL EHFAELFDLGFPNPLAERIIFETHQIHFANCSLVQPTFSDPPEDVLLAMIIAPI CLIPFLITLVVWRSKRTDTLL |
| pKB020 (R3wR2 C-tail) | METGALRRPQLLPLLLLLCGGCPRAGGCNETGMLERLPLCGKAFADMMG KVDVWKWCNLSEFIVYYESFTNCTEMEANVVGCYWPNPLAQGFITGIHRQ FFSNCTVDRVHLEDPPDEVLIPLIVIPVVLTVAMAGLVVWRSKRTDTLL |

### Table S6. Amino acid sequences of peptides

| | |
|---|---|
| CGRP(1-37) | ACDTATCVTHRLAGLLSRSGGVVKNNFVPTNVGSKAF-$NH_2$ |
| AM(13-52) | SFGCRFGTCTVQKLAHQIYQFTDKDKDNVAPRSKISPQGY-$NH_2$ |
| AM2/IMD(1-47) | TQAQLLRVGCVLGTCQVQNLSHRLWQLMGPAGRQDSAPVDPSSPHSY-$NH_2$ |
| CGRP(8-37) N31D/S34P/K35W/A36S | VTHRLAGLLSRSGGVVKNNFVPTDVGPWSF-$NH_2$ |
| AM(22-52) S48G/Q50W | TVQKLAHQIYQFTDKDKDNVAPRSKIGPWGY-$NH_2$ |
| AM(13-33)-AM2/IMD(28-47) "AM-AM2 half" | SFGCRFGTCTVQKLAHQIYQFMGPAGRQDSAPVDPSSPHSY-$NH_2$ |
| AM2/IMD(8-27)-AM(34-52) "AM2-AM half" | VGCVLGTCQVQNLSHRLWQLTDKDKDNVAPRSKISPQGY-$NH_2$ |
| AM(13-38)-AM2/IMD(34-47) "AM-AM2 ECD" | SFGCRFGTCTVQKLAHQIYQFTDKDKQDSAPVDPSSPHSY-$NH_2$ |
| AM2/IMD(8-33)-AM(39-52) "AM2-AM ECD" | VGCVLGTCQVQNLSHRLWQLMGPAGRDNVAPRSKISPQGY-$NH_2$ |
| AM(13-38)-AM2/IMD(23-33)-AM(39-52) "AM-AM2-AM" | SFGCRFGTCTVQKLAHRLWQLMGPAGRDNVAPRSKISPQGY-$NH_2$ |
| AM2/IMD(8-22)-AM(29-38)-AM2/IMD(34-47) "AM2-AM-AM2" | VGCVLGTCQVQNLSHQIYQFTDKDKQDSAPVDPSSPHSY-$NH_2$ |
| AM(13-52) N40K-TAMRA "AM-TAMRA" | SFGCRFGTCTVQKLAHQIYQFTDKDKD(K-TAMRA)VAPRSKISPQGY-$NH_2$ |
| AM2/IMD(8-47) D35K-TAMRA "AM2/IMD-TAMRA" | VGCVLGTCQVQNLSHRLWQLMGPAGRQ(K-TAMRA)SAPVDPSSPHSY-$NH_2$ |
